# Biocontainment strategies for *in vivo* applications of *Saccharomyces boulardii*

**DOI:** 10.1101/2022.12.27.522029

**Authors:** Karl Alex Hedin, Vibeke Kruse, Ruben Vazquez-Uribe, Morten Otto Alexander Sommer

**Affiliations:** Novo Nordisk Foundation Center for Biosustainability, Technical University of Denmark, 2800, Kgs. Lyngby, Denmark

**Author notes:** Corresponding authors, Morten Otto Alexander Sommer, Ruben Vazquez-Uribe.

**Keywords:** *S. boulardii*, Biocontainment, Biosafety, Probiotic yeast, Engineered microbes, Gut microbiome

## Abstract

The human gastrointestinal tract is a complex and dynamic environment, playing a crucial role in human health. Microorganisms engineered to express a therapeutic activity have emerged as a novel modality to manage numerous diseases. Such advanced microbiome therapeutics (AMTs) must be contained within the treated individual. Hence safe and robust biocontainment strategies are required to prevent the proliferation of microbes outside the treated individual. Here we present the first biocontainment strategy for a probiotic yeast, demonstrating a multilayered strategy combining an auxotrophic and environmental-sensitive strategy. We knocked out the genes *THI6* and *BTS1*, causing thiamine auxotrophy and increased sensitivity to cold, respectively. The biocontained *Saccharomyces boulardii* was unable to grow in the absence of thiamine above 1 ng/mL and exhibited a severe growth defect at temperatures below 20°C. The biocontained strain was well tolerated and viable in mice and demonstrated equal efficiency in peptide production as the ancestral non-biocontained strain. In combination, the data support that *thi6*Δ and *bts1*Δ enable biocontainment of *S. boulardii*, which could be a relevant chassis for future yeast-based AMTs.

## Introduction

The therapeutic potential of the human microbiome has gained significant attention, as microbiome-host interactions play a crucial role in various diseases (Canfora et al., 2019; Eckburg & Relman, 2007; Itzhaki et al., 2016; Sears & Garrett, 2014). Accordingly, the use of engineered living microbes to treat diseases is an emerging approach in the field of synthetic biology. Advanced microbiome therapeutics (AMTs) comprise microorganisms which have been genetically modified to express a therapeutic activity while present in the human microbiota. Previous work has demonstrated the use of AMTs to deliver therapeutic biomolecules (Arora et al., 2016; Chen et al., 2014; Steidler et al., 2000) or degrade toxic compounds (Isabella et al., 2018; Kurtz et al., 2019) in the gastrointestinal tract. The probiotic yeast *Saccharomyces boulardii* has recently caught attention as an AMT chassis, as it allows for the biosynthesis, folding and post-translational modification of several therapeutically relevant peptides and proteins (Nielsen, 2019). *S. boulardii* is a safe microorganism with over 40 years of use as a human probiotic (Kelesidis & Pothoulakis, 2012). However, an essential consideration in the design of AMTs is biocontainment strategies to limit the risk of the genetically modified microorganism proliferating outside the treated individual. Therefore, strategies for biocontainment must be developed for *S. boulardii* to ensure its future use as an AMT.

Effective biocontainment strategies should therefore take into consideration factors including mutagenetic drift, environmental supplementation, and horizontal gene transfer (Moe-Behrens et al., 2013). Several biocontainment strategies have been explored to prevent the proliferation and survival of engineered microorganisms in undesirable environments (Wook Lee et al., 2018). These strategies include auxotrophy (Bahey-El-Din et al., 2010; Isabella et al., 2018; Steidler et al., 2003), synthetic auxotrophy (Marlière et al., 2011; Pinheiro et al., 2012; Rovner et al., 2015), multispecies consortia (Johns et al., 2016; Mee et al., 2014), synthetic gene circuits (Chan et al., 2016; Huang et al., 2016; Piraner et al., 2016; Stirling et al., 2017), CRISPR-based kill switches (Caliando & Voigt, 2015; Rottinghaus et al., 2022), or a combination of them (Gallagher et al., 2015); however, each approach carries a risk. The auxotroph can be circumvented by scavenging the essential metabolite from nearby decayed cells or crossfeeding from established ecological niches. Synthetic auxotrophy may overcome these hurdles; however, it requires the gastrointestinal tract to be supplemented with the additional survival factor. Implementing synthetic gene circuits and CRISPR-based biocontainment strategies can lead to reduced fitness of the biocontained strain (Moe-Behrens et al., 2013), causing selective pressures to escape mutants (Uribe et al., 2021). Thus, a good biocontainment strategy requires a combination of strategies to compensate for each disadvantage.

While methods for biocontainment of bacterial AMTs have rapidly advanced (Bahey-El-Din et al., 2010; Isabella et al., 2018; Rottinghaus et al., 2022; Steidler et al., 2003), strategies for eukaryotic AMTs are still missing. In this study, we sought to develop a biocontainment strategy for *S. boulardii*. Since *S. boulardii* is a eukaryote, the risk of horizontal gene transfer is low (Emamalipour et al., 2020), and *S. boulardii* is unable to mate due to its sporulation defect (Lynne V. McFarland, 1996; Offei et al., 2019). Together, these factors reduce the risk of introducing genetic circuits or engineered DNA into natural ecosystems. Still, the risk of engineered *S. boulardii* proliferating in environments outside the treated host must be addressed. To address this issue, we sought to implement a biocontainment strategy for *S. boulardii* by reducing the fitness of the probiotic yeast outside the human host (Figure 1). We decided to evaluate the biocontainment capacity of both cold-sensitive and auxotrophic *S. boulardii* strains.

**Figure 1.**
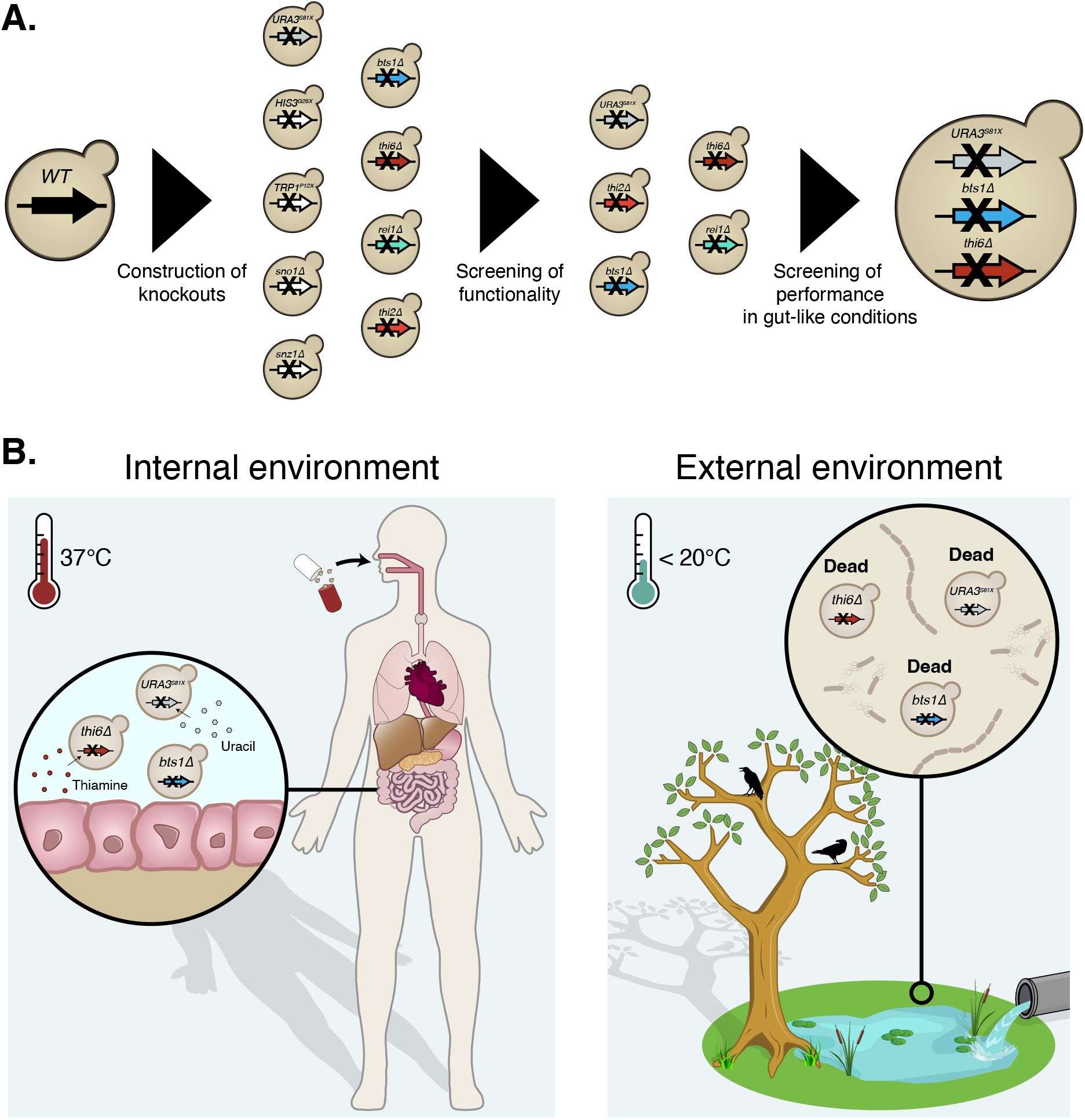
Graphical abstract of the implemented biocontainment strategy. **(A)** Schematic overview of the selection of the biocontainment strain. **(B)** Yeast cells with disruption of the genes synthesising uracil (*URA3*), thiamine (*THI6*), and geranylgeranyl diphosphate synthase (*BTS1*) can grow in the internal environment containing the essential nutrients and optimal temperature. However, proliferation in external environments lacking the essential nutrients and lower temperatures will be limited.

## Methods

### Plasmids and strain construction

This study’s primers, plasmids, gBlocks and gRNA sequences are listed in Supplementary Table S1, S2, S3, and S4. Oligonucleotides and gBlocks were ordered from Integrated DNA Technologies (IDT). To generate the *URA3^S81X^* (SbU^-^), *HIS3^G26X^* (SbH^-^), and *TRP1^P12X^* (SbT^-^) disrupted strains, gBlocks with respectively gRNA were assembled with pCfB2312 (Cas9-KanMX) and transformed with their respective donor primer (Supplementary Table S1). To generate the *thi2*Δ, *thi6*Δ, *snz1*Δ, *sno1*Δ, *rei1*Δ and *bts1*Δ strains, gBlocks from respectively gRNA were assembled with pCfB3050 (pCas9-*URA3*) and transformed with respectively donor primer (Supplementary Table S1). The GFP, RFP and Exendin-4 integration plasmid (Supplementary Table S2) was generated by assembly of respective gBlocks (Supplementary Table S3) with pCfB2909 (marker free) and transformed together with pCfB6920.

All plasmid assemblies were conducted with Gibson Assembly (Gibson et al., 2009) and transformed into One Shot^®^ TOP10 *Escherichia coli* (Thermo Fisher Scientific). All *E. coli* were grown in lysogeny broth (LB) media containing 5 g/L yeast extract, 10 g/L tryptone and 10 g/L NaCl; (Sigma Aldrich) supplemented with 100 mg/L ampicillin sodium salt (Sigma Aldrich). LB agar plates contained 1% agar (Sigma Aldrich).

*S. boulardii* (*S. cerevisiae* ATCC^®^ MYA796^™^) was obtained from American Type Culture Collection (ATCC). The strains created in this study are listed in Supplementary Table S5. *S. boulardii* transformations were performed via high-efficiency yeast transformation using the LiAc/SS carrier DNA/PEG method (Gietz & Woods, 2006). Genomic integrations cassettes were digested with the restriction enzyme NotI (FastDgiest Enzyme, Thermo Scientific^™^) prior to transformation and transformed together with various helper plasmids and pre-expressed Cas9 from pCfB2312. All transformations were incubated at 30°C for 30 minutes and then heat-shocked in a water bath at 42°C for 60 minutes. All transformations followed a recovery step. The transformation tubes were microcentrifuge for 2 min at 3000 g. Pellets were resuspended in 500 μL of YPD media containing 10 g/L yeast extract, 20 g/L casein peptone and 20 g/L glucose (Sigma Aldrich) and incubated for 2 to 3 hours at 30°C before being plated. All yeast transformations were plated on synthetic complete (SC) plates containing 1.7 g/L yeast nitrogen base without amino acids and ammonium sulphate (Sigma Aldrich), 1 g/L monosodium glutamate (Sigma Aldrich), 1.92 g/L Yeast Synthetic Drop-out Medium

Supplements without uracil (Sigma Aldrich) and 200 mg/L geneticin (G418; Sigma Aldrich) at 37°C. Colony-PCR using OneTaq (Thermo Scientific^™^) confirmed the genomic integration. Primers flanking the integration were used to confirm the integration. Genomic DNA was extracted by boiling cells at 70°C for 30 min in 20 mM NaOH. The strains were cured for pCfB2312 and helper plasmids after genome integration.

### Cultivation experiments

All cultivation was started from a pre-culture, inoculated with a recently streaked out colony from a −80°C cryostock, cultivated for 12 – 16 hours. All pre-cultures were conducted in the same media as the following cultivation experiment unless other stated. Pre-culture to determine the auxotrophic strains was done in media supplemented with the required nutrition and washed three times in 1% PBS to ensure no traces of the required nutrition were transferred into the new cultures. Pre-culture to determine cold sensitivity was conducted at 37°C to ensure enough biomass.

### Real-time growth monitoring

Real-time OD_600_ measurements were obtained every 10 min for approximately 72 hours with microplate reader Synergy^™^ H1 (BioTek) for aerobic and microaerobic conditions. Epoch 2 microplate reader (BioTek) was used for anaerobic real-timer OD_600_ measurements. All cultures had an initial OD_600_ of 0.05. The cultures were incubated into 200 μL minimal synthetic complete media (DELFT) containing 7.5 g/L (NH_4_)_2_SO_4_, 14.4 g/L KH_2_PO_4_, 0.5 g/L MgSO_4_ x 7H_2_O, 20 g/L glucose, 2 mL/L trace metals solution, and 1 mL/L vitamins with or without thiamine and pyridoxine (Verduyn et al., 1992), and with or without 20 mg/L uracil, 20 mg/L histidine and 20 mg/L tryptophan supplemented. CELLSTAR^®^ 96 well cell culture plate (Greiner Bio-One) with an air-penetrable lid (Breathe-Easy, Diversified Biotech) was used for all cultivation. pH was adjusted with 1M HCl to 3, 4, 5 and 6 for the respective experiment. Cultivation was performed with continuous double orbital shaking of 548 cycles per minute (CPM) at 37°C and 0%, 0.1%, 1% or 21% oxygen. Anaerobic conditions were obtained using a vinyl anaerobic chambers (Coy Laboratory Products Vinyl; gas mixture, 95% N_2_ and 5% H_2_), and microaerobic conditions were obtained using CO_2_/O_2_ Gas Controller (BioTek).

### Thiamine dose-response experiment

The cultures were incubated into 200 μL SC media without thiamine, containing 1.7 g/L yeast nitrogen base without amino acids and thiamine (FORMEDIUM^™^), 1.92 g/L Yeast Synthetic Drop-out Medium Supplements without uracil, for 48 hours with an initial OD_600_ of 0.05. The media was supplemented with either 0, 0.001, 0.01, 0.1, 1, or 400 μg/mL thiamine hydrochloride (Sigma Aldrich) and 20 mg/L uracil.

### Escape rate experiment of the thiamine auxotroph

The SbU^-^ and SbU^-^ +*thi6*Δ strains were incubated in 20 mL YPD for 72 hours. The cultures were incubated in a 250 mL shake flask with an initial OD_600_ of 0.05. The culture was spun down and diluted in MilliQ water to OD_600_ 10. A serial dilution from OD_600_ 10 to 0.00001 was generated. 100 μL of the undiluted and 5 μL from each dilution were plated on SC plates with and without thiamine supplemented (Supplementary Figure S2B). The plates were incubated at 37°C for 72 hours. Cell mass from the undiluted 5 μL spotting was spread out on SC plates with and without thiamine supplemented (Supplementary Figure S2C).

### Cold exposure experiment

The SbU^-^, SbU^-^ +*rei1*Δ, SbU^-^ +*bts1*Δ and SbU^-^ +*bts1*Δ + *thi6*Δ strains were incubated in 2.6 mL YPD in a 24-deep well plate (Axygen^®^, VWR) with a sandwich cover (Enzyscreen) and with an initial OD_600_ of 0.05. The plates were incubated at 15, 20 and 37°C for a maximum of 120 hours. OD_600_ was measured at 0, 8, 24, 32, 48, 72, 96, and 120 hours.

### Competition experiment in the presence and absence of thiamine

Single and co-cultivation of (SbU^-^)-GFP and (SbU^-^ +*bts1*Δ + *thi6*Δ)-mKate were cultured in 2 mL SC media with and without thiamine in a 24-deep well plate (Axygen^®^, VWR) with a sandwich cover (Enzyscreen) and with an initial OD_600_ of 0.1. The co-culture was started with a 1:1 ratio of each strain. 20 μL of single and co-cultures were transferred into fresh media every 48^th^ hours for a total period of 96 hours (two transfers).

### Competition experiments in different temperature

Single and co-cultivation of (SbU^-^)-GFP and (SbU^-^ + *bts1*Δ + *thi6*Δ)-mKate were cultured in 2 mL SC media YPD in a 24-deep well plate (Axygen^®^, VWR) with a sandwich cover (Enzyscreen) and with an initial OD_600_ of 0.1 at 15°C, 20°C, and 37°C. The co-culture was started with a 1:1 ratio of each strain. 20 μL of single and co-cultures were transferred into fresh media every 24^th^ hours for a total period of 120 hours (five transfers) for the 20°C and 37°C cultivation. Single and co-cultures at 15°C were transferred into fresh media every 48^th^ hours for a total period of 144 hours (three transfers). The culture was transferred every 48^th^ hours to ensure that SbU^-^ reached similar cell counts at all temperatures (Supplement Figure S6).

### Survival assay

Serial dilution (1,000x and 10,000x) from the 37°C co-cultivation of (SbU^-^)-GFP and (SbU^-^ +*bts1*Δ + *thi6*Δ)-mKate in YPD were plated on YPD plates and incubated at 15°C, 20°C and 37°C for 144, 96, and 48 hours respectively. Red and green colony forming units (CFU) were counted under blue light.

### Flow cytometry

All competition experiments were analysed with a NovoCyte Quanteon^™^ (Agilent) flow cytometry. 20 μL culture was diluted in 180 μL 1% PBS and run on the flow cytometry with a threshold of 5,000 yeast events or 60 μL sample injection. Yeast events were gated based on size events (FSC-A) < 6 x 10^6^ and complexity events (SSC-A) < 2 x 10^5^. Singlets were gate based on SSC (SSC-A vs SSC-H). The (SbU-)-GFP population was quantified with the FITC channel, and (SbU^-^ +*bts1*Δ + *thi6*Δ)-mKate was quantified with PE-TexasRed. The population distribution was quantified based on FITC-H vs PE-TexasRed-H. Absolute events were calculated.

### ELISA

The Exendin-4 producing *S. boulardii* strains were incubated in 2 mL DELFT medium supplemented with 20 mg/L uracil in a 24-deep well plate (Axygen^®^, VWR) with a sandwich cover (Enzyscreen). The cultures had an initial OD_600_ of 0.05 and were performed with continuous shaking at 250 RPM at 37°C. All cultures were harvested after 24 and 48 hours. Cell cultures were spun down at 10,000 g for 10 min at 4°C. Exendin-4 was quantified with Exendin-4 EIA (EK-070-94, Phoenix). The signals were detected by OD_450_ using a microplate reader Synergy^™^ H1 (BioTek).

### Animal experiment

All animal experiments were conducted according to the Danish guidelines for experimental animal welfare, and the study protocols were approved by the Danish Animal Experiment Inspectorate (license number 2020-15-0201-00405). The study was carried out in accordance with the ARRIVE guidelines (du Sert et al., 2020). All *in vivo* experiments were conducted on male C57BL/6 mice (6-8 weeks old; Taconic Bioscience). Unless otherwise stated, all mice were housed at room temperature on a 12-hour light/dark cycle and given *ad libitum* access to water and standard chow (Safe Diets, A30). All mice were randomised according to body weight and acclimated for at least one week prior to the first oral administration. Each animal study received a freshly prepared batch of *S. boulardii*. Body weight and food intake were recorded once per week. The researchers were blinded in all mouse experiments. The mice were euthanised by cervical dislocation at the end of the study.

The mice were divided into four groups (n = 4), either receiving the Sb wild-type, Sb *bts1*Δ, Sb *thi6*Δ or Sb *bts1*Δ + *thi6*Δ strain. The mice were orally administered via intragastric gavage with ~10^8^ CFU of the *S. boulardii* strain in 100 μL of 1x PBS and 20% glycerol. The mice were orally administered with *S. boulardii* for five consecutive days, followed by a six-day washout. The drinking water was supplemented with an antibiotic cocktail containing 0.3 g/L ampicillin sodium salt, 0.3 g/L kanamycin sulfate, 0.3 g/L metronidazole, and 0.15 g/L vancomycin hydrochloride after the washout period. After five days of antibiotic treatment, the mice were orally administered with *S. boulardii* in 100 μL of 1x PBS (containing 1.0 g/L ampicillin sodium salt, 1.0 g/L kanamycin sulfate, 1.0 g/L metronidazole, and 1.0 g/L vancomycin hydrochloride) and 10% glycerol for five consecutive days. The washout for the antibiotic-treated mice was monitored for 33 days.

The faeces were collected in pre-weighed 1.5 mL or 2.0 mL Eppendorf tubes containing 1 mL of 1x PBS and 50% glycerol and weighed again to determine the faecal weight. All sample preparation for assessing CFU numbers was kept on ice and followed the same practice. The faecal samples were homogenised by vortexed at ~2400 rpm for 20 min. The samples were then spun down at 100 g for 30 seconds, followed by a dilution series, where 5 μL of each dilution was plated in duplicates or triplicates. Under the antibiotic-treated washout period, 100 μL was plated. The faecal samples were plated on SC supplemented with 20 mg/L uracil plates containing 100 mg/L ampicillin, 50 mg/L kanamycin, and 30 mg/L chloramphenicol (Sigma Aldrich).

### Statistical testing

All statistical analysing were performed in RStudio version 4.1.0 with the rstatix and DescTools packages. Unless otherwise stated, all data are presented as means + SEM. Statistical differences between groups of two were analysed with a dependent sample t-test. Bonferroni adjustments were used for multiple comparisons. Comparison of three or more groups was analysed by ANOVA with either Tukey’s HSD post hoc test or Dunnett’s post hoc test. The statistical significance level was set at p < 0.05.

## Results

### Selection of optimal auxotrophies for *S. boulardii* biocontainment

We started by investigating the impact of constraining the probiotic yeast to become dependent on the exogenous supply of metabolites for growth and survival, as it is one of the most common biocontainment strategies for genetically modified microorganisms (Wook Lee et al., 2018). Here we generated a library of auxotrophic *S. boulardii* strains by disrupting several genes and evaluating the burden of the gene deletion on each strain. We created one nucleoside (uracil), two amino acids (histidine and tryptophan) and two vitamins (thiamine and pyridoxine) auxotrophic strains (Figure 2A). We started by introducing a stop codon in *URA3* (uracil synthesis), *HIS3* (histidine synthesis) and *TRP1* (tryptophan synthesis), as these auxotrophic strains have previously been reported in *S. boulardii* (Liu et al., 2016). Importantly, these gene disrupted strains are suitable hosts for further genetic manipulation using existing *Saccharomyces cerevisiae* tools (Durmusoglu et al., 2021; Hudson et al., 2014; Jin et al., 2021; Liu et al., 2016). The strains with *URA3, HIS3*, and *TRP1* disrupted were unable to grow unless uracil, histidine or tryptophan was supplemented to the growth medium (Figure 2B-C). The *TRP1^P12X^* (SbT^-^) strain also showed a metabolic burden when media was supplemented with tryptophan (Figure 2C, Supplementary Figure S1). Based on these observations, the *HIS3^G26X^* (SbH^-^) and *URA3^S81X^* (SbU^-^) strains showed to be the most promising nutritional auxotrophs compared to the SbT^-^ strain. The SbU^-^ strain was selected for further gene knockouts based on a previous report showing higher gene expression from *URA3-* marker plasmids (Durmusoglu et al., 2021).

**Figure 2.**
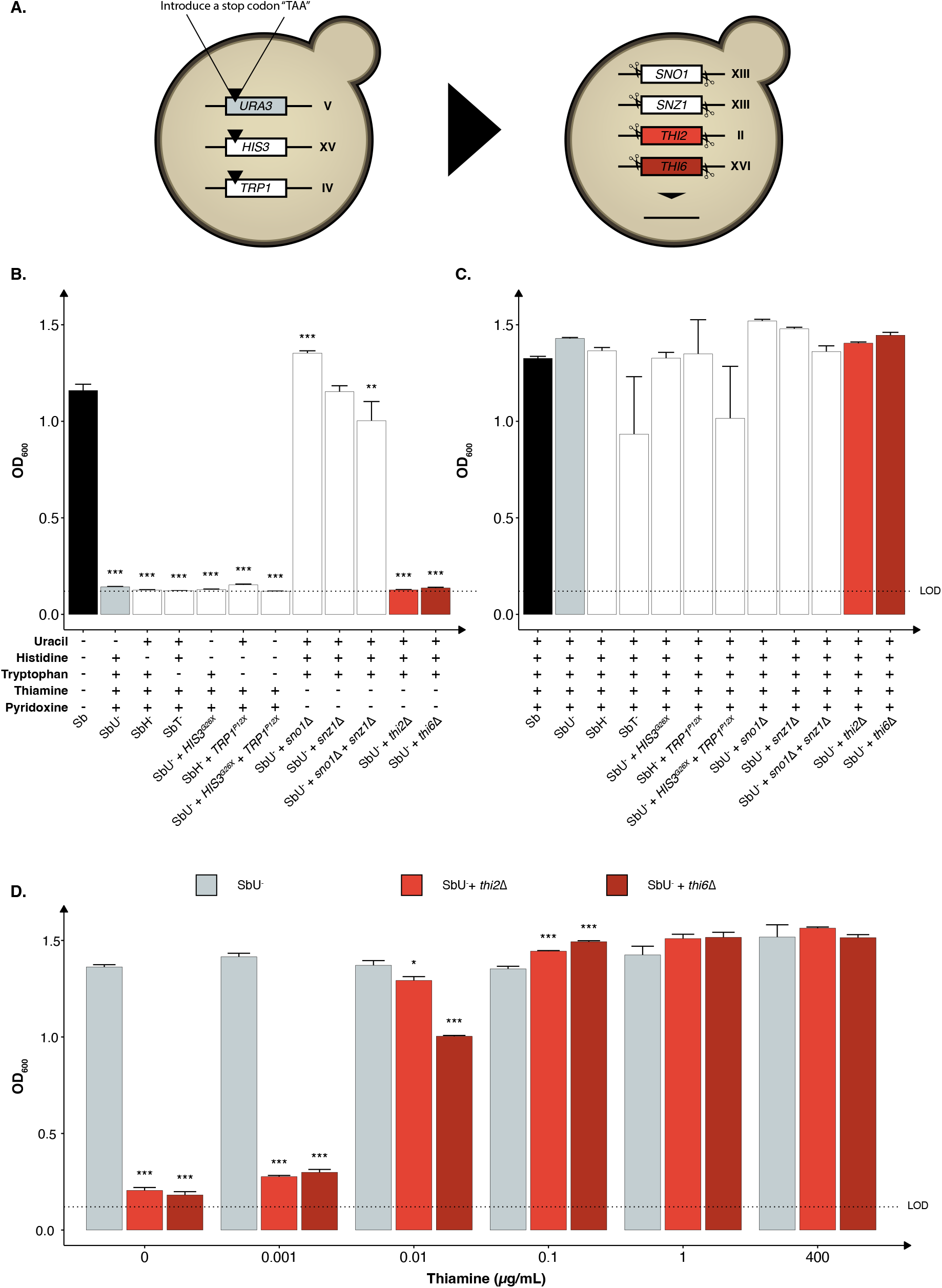
Selection of auxotrophic mutation in *S. boulardii*. **(A)** Schematic overview of the construction of the auxotrophic strains. To generate the uracil, histidine, and tryptophan auxotroph, a null mutation was introduced in the respective genes, while the genes were deleted to generate the vitamin auxotrophs. Bar plot of mean OD_600_ after 48 hours **(B)** without the required nutrition supplemented and **(C)** with the required nutrition supplemented. **(D)** Bar plot of mean OD_600_ after 48 hours under different concentrations of thiamine. Limit of detection (LOD). Data presented as mean +SEM (n = 3). * p < 0.05, ** p < 0.01 and *** p < 0.001. One-way ANOVA, Dunnett’s post hoc test with Sb **(A)** or SbU^-^ **(B)-(C)** as reference.

We further investigated the phenotype of disrupting two other biosynthetic pathways in strains with uracil synthesis deficiency to determine potential additive effects. Knocking out either the *THI2* or the *THI6* gene in the thiamine synthesis pathway resulted in strains that were unable to grow in media lacking thiamine (Figure 2B). Knocking out the *SNO1* and *SNZ1* genes of the pyridoxine synthesis pathway individually resulted in no growth defect, contrary to previously reported (Rodríguez-Navarro et al., 2002). Nonetheless, a significant growth defect was observed when knocking out the *SNO1* and *SNZ1* genes in combination. To demonstrate that the growth defect resulted from the respective auxotrophies, we cultured the various strains in media containing the corresponding nutrient supplements and observed similar OD_600_ reaching after 48 hours (Figure 2C).

To examine the robustness of the auxotrophic strains towards conditions experienced in the human gastrointestinal tract, we assessed their growth performance in minimal media with pH and oxygen conditions more closely mimicking the gastrointestinal tract. The *thi2*Δ strain displayed a more considerable metabolic burden compared to *thi6*Δ, demonstrating a growth defect at pH 6 and in microaerobic (0.1% oxygen) and anaerobic conditions (Supplementary Figure S1). The *thi6*Δ strain showed a slight growth defect at pH 4 (Supplementary Figure S1) while generally performing better than the *thi2*Δ strain.

We next evaluated the thiamine sensitivity in the strains to identify the minimum thiamine concentration required for growth by the *thi2*Δ and *thi6*Δ strains to circumvent the biocontainment. Both knockouts showed a growth defect at a concentration < 0.1 μg/mL (Figure 2D). The *thi6*Δ strain appeared more sensitive to low thiamine concentrations than the *thi2*Δ strain. We also confirmed that from a 72-hour culture, no escapers were identified for the *thi6*Δ strain (Supplementary Figure S2).

### Constructing cold-sensitive *S. boulardii* strains

Another strategy to reduce the proliferation of genetically modified microorganisms in specific environments is to make them sensitive to temperature changes that may occur outside the targeted environment (Figure 3A). To assess this approach, we knocked out two genes (*REI1* and *BTS1*) that were previously reported to exhibit a growth defect at a temperature lower than 25°C when knocked out in *S. cerevisiae* (Hung & Johnson, 2006; Jiang et al., 1995; Lebreton et al., 2006a). The *REI1* gene encodes a zinc finger protein part of the 60S complex, and the *BTS1* gene encodes the yeast geranylgeranyl diphosphate synthase. We observed the expected growth defects at temperatures of 15°C and 20°C for both *rei1*Δ and *bts1*Δ strains (Figure 3B). The *rei1*Δ strain displayed a more pronounced growth defect, showing no growth at 15°C after 120 hours. At 20°C, the *rei1*Δ strain started growing after 72 hours (Supplementary Figures S3). Both *rei1*Δ and *bts1*Δ strains were hypersensitive at 15°C; however, the *rei1*Δ strain also showed a growth defect at 37°C (Supplementary Figure S3).

**Figure 3.**
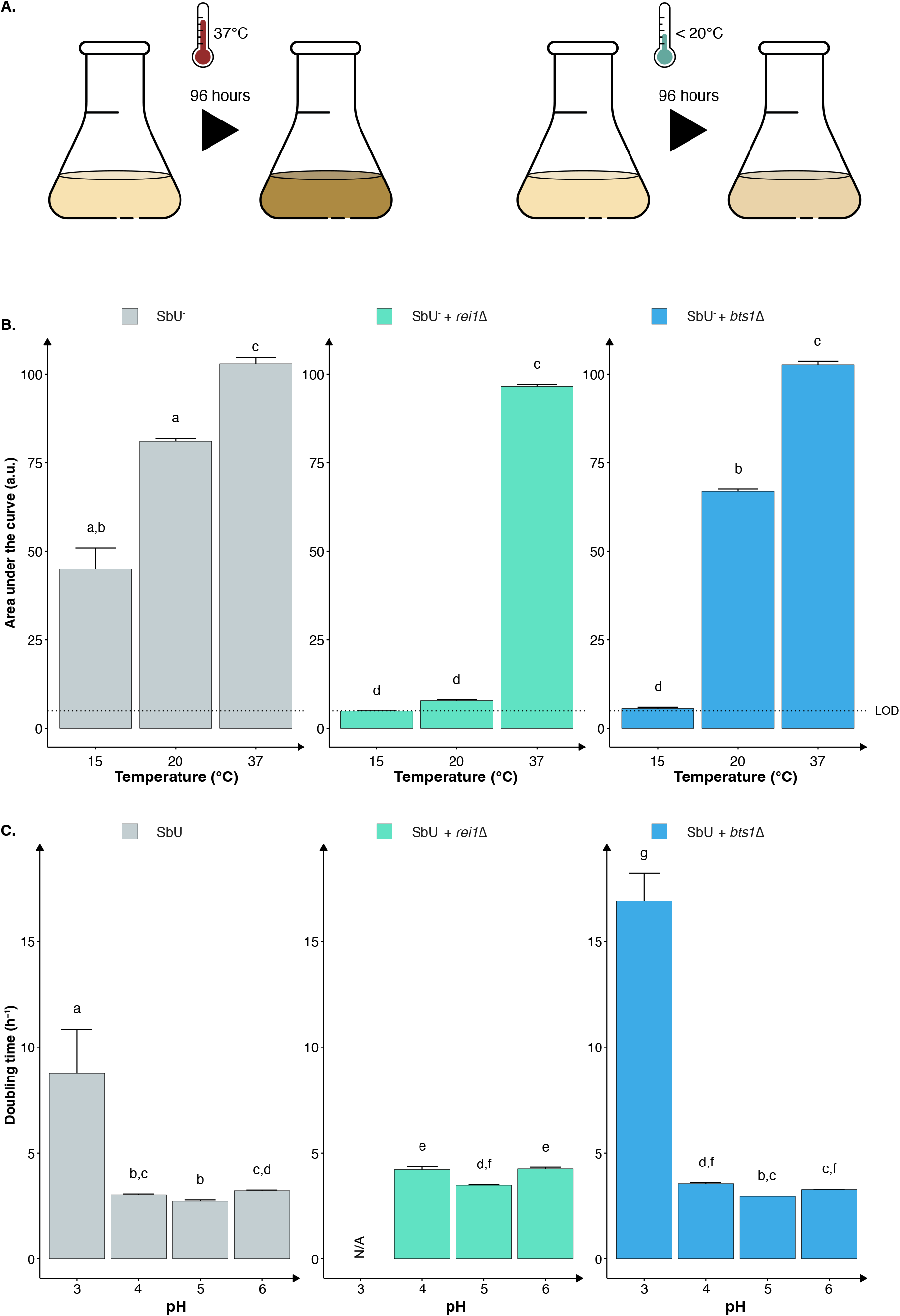
Constructing temperature-sensitive *S. boulardii* strains. **(A)** Graphical illustration of the experimental design to confirm the phenotype. **(B)** Bar plot of the mean area under the curve (AUC) of 96-hour cultivation at 15°C, 20°C and 37°C. **(C)** Bar plot of the mean doubling time in pH 3, 4, 5 and 6. Limit of detection (LOD). Data presented as mean + SEM (n = 3). Two-way ANOVA, Tukey post hoc test. The different letters (a, b, c, d, e, f, g, and h) above the bars indicate statistically different groups (significance level at p < 0.05)

Furthermore, evaluating the two cold-sensitive strains in the gut-like conditions, we demonstrate that the *rei1*Δ strain showed a more pronounced fitness cost in minimal media at different pH (Figure 3C). The *rei1*Δ strain was unable to grow at a pH lower than 4 after 72 hours, and the strain displayed an approximate 25% slower doubling time at pH 4, 5, and 6 compared to the parental strain (SbU^-^). The *rei1*Δ strain also showed a significantly lower growth rate in anaerobic and microaerobic conditions than SbU^-^ (Supplementary Figure S4), while the *bts1*Δ demonstrated a slower growth rate in anaerobic conditions. The growth defect at anaerobic conditions was comparatively more pronounced for the *bts1*Δ than for the *rei1*Δ strain.

### Combining vitamin auxotrophy with cold-sensitive mutations as a robust biocontainment strategy

To generate a more robust biocontainment strain, we combined the nutritional auxotrophy and the cold-sensitive mutant (Figure 4A). Specifically, we chose *thi6*Δ and *bts1*Δ, in the SbU^-^ background strain, based on functionality at different pH concentrations and oxygen gradients. The double gene knockout of *BTS1* and *THI6* maintained the phenotypic effect of the individual knockouts, demonstrating the inability to grow in the absence of thiamine supplementation and slower growth in temperatures lower than 20°C (Supplementary Figure S3 and Supplementary Figure S5).

**Figure 4.**
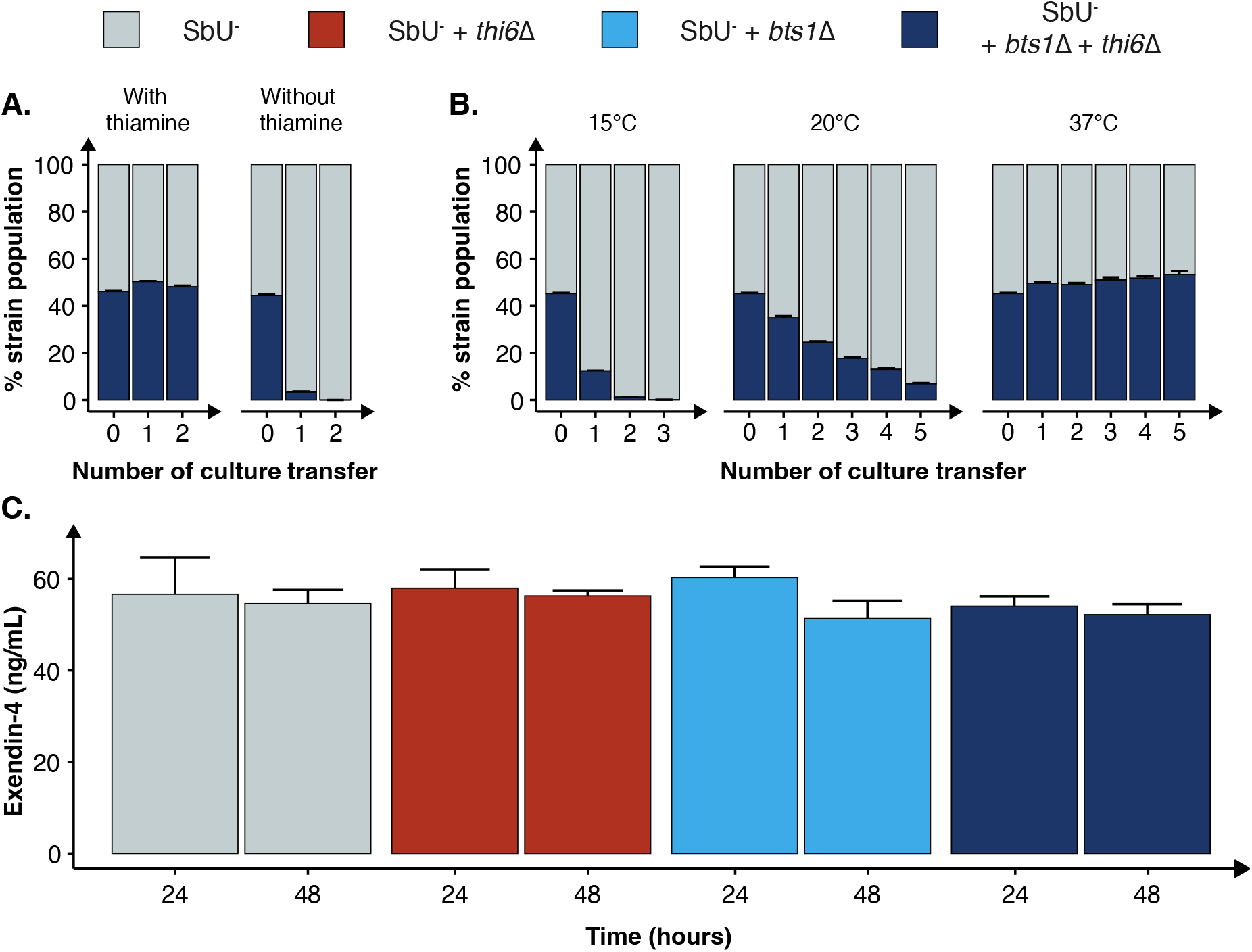
Characterisation of the combined cold-sensitive and auxotrophic strain. **(A)** Stacked bar plot of the percentage of SbU^-^ and SbU^-^ + *bts1*Δ +*thi6*Δ in a co-culture experiment with and without thiamine. The co-culture was transferred to a fresh culture every 48 hours (n = 5). **(B)** Stacked bar plot of the % of SbU^-^ and SbU^-^ + *bts1*Δ + *thi6*Δ in a co-culture experiment at 15°C, 20°C and 37°C. The co-culture was transferred to a fresh culture every 24 or 48 hours (n = 5). **(C)** Bar plot of the mean concentration of Exendin-4 (ng/mL) quantified in the supernatant at 24 and 48 hours of cultivation (n = 3). Data presented as mean + SEM. * p < 0.05, One-way ANOVA, Dunnett’s post hoc test with SbU^-^as reference.

Next, we evaluated the competitive fitness of the double knockout biocontainment strain in a co-culture with the parental control strain SbU^-^ in the presence and absence of thiamine and at different temperatures. The double knockout and control strain perform similarly in the presence of thiamine; however, in the absence of thiamine, the double knockout strain was undetected after two transfers (Figure 4B and Supplemented Figure S6). In the same manner, the double knockout strain and the control strain were able to grow simultaneously at the permissive temperature conditions of 37°C; however, at 15°C and 20°C, the percentage of double knockout strains was drastically reduced incrementally with the number of culture transfers (Figure 4C). The biocontainment double knockout strain maintained ~1% of the total yeast population after two transfers of the co-culture at 15°C and ~5% after five transfers at 20°C.

Furthermore, we tested whether the biocontainment strain would maintain its cold-sensitive phenotype after multiple generations in co-culture with the control strain at 37°C (120 hours). Here, we observed that the double knockout strain maintained its cold-sensitive phenotype, demonstrating zero colonies after plating the culture, incubating the strain at 15°C and maintaining a slower growth phenotype at 20°C (Supplementary Figure S7). The double knockout biocontainment strain also showed no synergistic growth defect at different pH concentrations and oxygen percentages (Supplementary Figure S8). We also observed an increased thiamine sensitivity by the combined biocontainment strain, showing a significantly lower OD_600_ in 1 μg/mL after 48 hours (Supplementary Figure S9).

Finally, we sought to investigate if the double knockout biocontainment strain can produce a similar level of a recombinant protein as the parental strain SbU^-^. This is relevant since some gene knockouts might negatively impact protein synthesis (Puddu et al., 2019). Therefore, we engineered the biocontainment strains to produce and secrete a GLP-1 receptor agonist (Exendin-4) since *in situ* biosynthesis of Exendin-4 in the gastrointestinal tract with *S. boulardii* has previously been shown to reduce food intake and body weight in mice (Hedin, Zhang, et al., 2022). We observed no significant difference in the production of Exendin-4 after 24 and 48 hours (Figure 4D), suggesting that the biocontainment strain does not compromise the production of a heterologous protein.

### The biocontainment strain is safe and viable in a mouse model

While we have validated the biocontainment strategy to be functional *in vitro*, the generated knockout strains still need to be demonstrated to be safe and viable in the gastrointestinal tract to express a therapeutic activity. To assess this, we orally administered the biocontainment strains daily for five days in conventional mice with an intact microbiome (Figure 5A) to study if the biocontainment strains pose any fitness cost while interacting with other microbes. We demonstrated that the wild-type, single and combined knockouts colonise equally in the mice with intact microbiome (Figure 5B). In addition, all the *S. boulardii* strains were washed out after six days, except for one mouse receiving the *bts1*Δ strain (Figure 5C), indicating that the knockouts did not cause any unwanted engraftment in mice.

**Figure 5.**
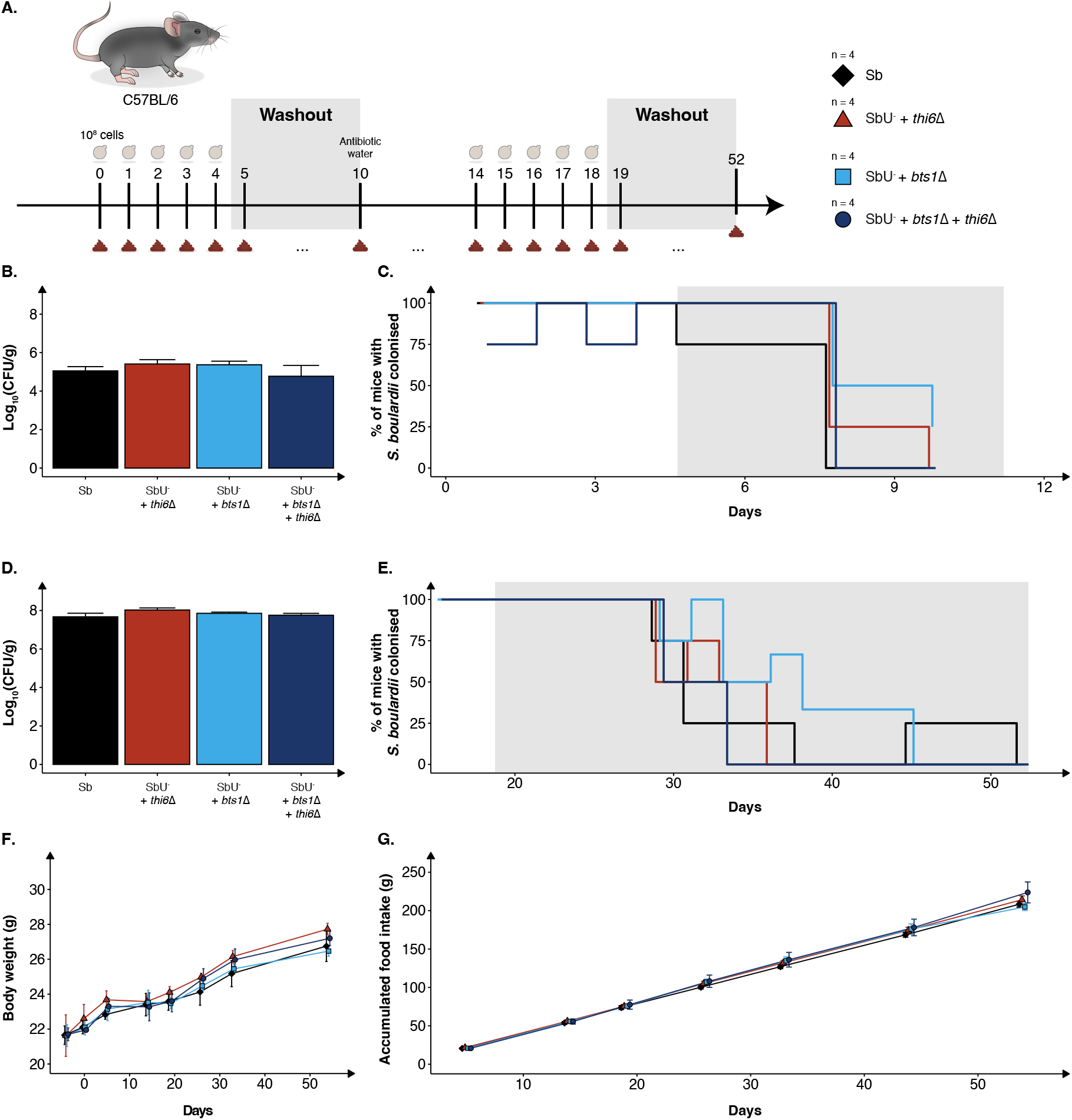
*In vivo* assessment of the biocontainment strains in antibiotic-treated mice. **(A)** Graphical scheme of the study design. Male C57BL/6 mice were orally administered with ~10^8^ cells of *S. boulardii* daily for five successive days, followed by six days of washout. On the 10^th^ day was, the water supplemented with an antibiotic cocktail. The mice were again orally administered with ~10^8^ cells of *S. boulardii* daily for five successive days, followed by 34 days of washout. **(B)** Bar plot of *S. boulardii* abundance (log_10_ CFU per gram faeces) in conventional mice with intact microbiome. **(C)** Step plot of percentage of conventional mice with *S. boulardii* colonised (LOD ≈ 10^3^ CFU/g). **(D)** Bar plot of *S. boulardii* abundance (log_10_ CFU per gram faeces) in antibiotic-treated mice. **(E)** Step plot of percentage of antibiotic-treated mice with *S. boulardii* colonised (LOD ≈ 50 CFU/g). **(F)** The body weight (grams) during the whole study. **(G)** Accumulated food intake (gram) throughout the whole study. Data presented as mean ± SEM (n = 4). Data were analysed with One-way ANOVA (B and D) and Twoway ANOVA (F and G), using Dunnett’s post hoc test with SbU^-^ as reference.

Furthermore, we evaluated the biocontainment strains in an antibiotic-treated mouse model designed for pre-clinical testing of yeast-based AMTs (Hedin, Rees, et al., 2022) and investigated if the strains could colonise the murine gastrointestinal tract. The antibiotic-treated mice were orally administered with the biocontainment strains for five days. Wild-type, single and combined knockouts were demonstrated to also colonise equally in the mice with a reduced microbiota (Figure 5D). Although the *S. boulardii* was washout slower in the antibiotic-treated mice, no mice displayed any detectable levels 33 days after the last oral administration (Figure 5E).

To investigate if the newly generated strains posed any safety concerns to the health and well-being of the mice, body weight, food intake and behaviour were monitored throughout the whole study. We observed no change in body weight or food intake in the mice receiving the biocontainment strains (Figure 5F-G), both during low and high colonisation of the strains. In addition, no abnormal behaviour was observed in the mice.

## Discussion

Biocontainment is a crucial step in the development of AMTs, as it constrains the proliferation of genetically modified microorganisms outside the treated individual. Therefore, it limits the risk of outcompeting natural organisms and negatively affecting ecosystems and human health (Wilson, 2008). In this study, we implemented a biocontainment strategy for the probiotic yeast *S. boulardii*, demonstrating robust growth in the murine gastrointestinal tract and limited growth under in vitro conditions mimicking the external environments (Figure 1).

We designed a multi-layered strategy by introducing a combination of auxotrophy and temperature sensitivity. We selected the SbU^-^as a background strain for further genetic manipulation as it was demonstrated to have the expected phenotypic traits as auxotroph and *URA3* marker plasmids have been reported to have higher gene expression than *HIS3* and *TRP1* marker plasmids in *S. boulardii* (Durmusoglu et al., 2021) and is a frequently used marker (Jensen et al., 2014; Maury et al., 2016). Next, we evaluated the effect of additional auxotrophic knockouts as potential strategies. While the *thi2*Δ and *thi6*Δ strains showed no growth in the functionality assay (Figure 2B), both showed a slight increase in OD_600_ at 0 μg/mL thiamine in the thiamine dose experiment (Figure 2D). This could be explained by the fact that a small trace amount of thiamine could be transferred from the pre-culture; alternatively, the strains are more sensitive in media lacking pyridoxine. Nonetheless, the thiamine auxotrophic strain, *thi6*Δ, demonstrated a more pronounced and sensitive effect to lower thiamine concentrations than *thi2*Δ. In addition, the *THI2* is a transcriptional activator of thiamine biosynthetic genes (Nishimura et al., 1992), while *THI6* is a gene encoding an enzyme required for thiamine synthesis (Nosakas et al., 1994); as such, circumventing the transcriptional activation might pose a higher risk of escapers than gaining back an enzymatic function.

To further contain the probiotic yeast, we investigated the potential of temperature-sensitive knockout strains. The global average temperature is calculated to be 13.9°C (Sobrino et al., 2020). Accordingly, an AMT with reduced fitness in that temperature range but a stable and robust growth at 37°C would be of interest. We compared two gene knockouts that caused the strain to be sensitive to colder temperatures (Hung & Johnson, 2006; Jiang et al., 1995). Knocking out *BTS1* and *REI1* reduced the fitness of the strains at temperatures ≤ 20°C. We prioritised the *bts1*Δ strain even though the *rei1*Δ strain showed a more pronounced growth defect at 20°C. This was based on the fact that the *rei1*Δ strain also displayed a slower growth at 37°C and an increased fitness cost in different pH and oxygen conditions found in the gastrointestinal tract. Furthermore, it has been reported that overexpression of *REH1* and deleting *ARX1* can partially suppress the *rei1*Δ cold-sensitive growth phenotype (Lebreton et al., 2006b; Parnell & Bass, 2009). Thus, the *rei1*Δ strain might increase the risk of selective pressures for escape mutants.

In addition, we combined the auxotrophic mutant *thi6*Δ and the cold-sensitive mutant *bts1*Δ in the background strain SbU^-^ to further reduce the fitness in the external environments and minimise the chance of escape mutants. The final biocontainment strain exhibited similar phenotypic traits as the individual knockouts. We also demonstrated that the parental control strain drastically outcompetes the double biocontainment strain unless thiamine was supplemented and cultivated at 37°C. Furthermore, when thiamine was absent from the media, the double biocontainment strain was equally restricted in mono-culture as in co-culture with the control strains, indicating any potential cross-feeding of thiamine between the strains was not sufficient to maintain the growth of the biocontainment strain.

The knockout strains demonstrated a neglectable effect on peptide synthesis, indicating that the biocontainment strain could still act as an AMT for delivering therapeutic peptides. Furthermore, we observed comparable phenotypic growth performance at the different pH and oxygen conditions, demonstrating the strain to be robust to potential environmental changes.

We finally determined the viability and safety profile of the single and double knockout strains in healthy mice. We observed no differences in the mice health and well-being between the different groups, demonstrating that the knockout strains did not pose a risk in healthy mice. The viability of the biocontainment strains in the faeces was demonstrated to reach equal levels compared to the wild-type, both in naïve and antibiotic-treated mice. The abundance per gram of faeces and washout of *S. boulardii* in mice’s gastrointestinal tract is in line with previously reported data (Hedin, Rees, et al., 2022). This supports the hypothesis that the biocontainment strain does not suffer any noticeable fitness cost in the gastrointestinal tract of a mouse. In addition, it demonstrates that the concentration of thiamine in the murine gut is sufficient for the *thi6*Δ strain to grow.

Moving AMTs into humans requires strategies that ensure the safety of the AMT chassis. Our study demonstrates a multi-layered biocontainment strategy validated both under laboratory growth conditions and *in vivo* in the gastrointestinal tract of mice. Although further combinations of strategies might be required to ensure complete containment of the yeast, we here demonstrate, to the author’s knowledge, the first biocontainment strategy as a platform for the continued development of yeast-based AMTs.

## Data availability

This published article and its supplementary material include all relevant data generated or analysed during this study. Further inquiries can be directed to the corresponding author/s.

## Acknowledgement

This work received funding from The Novo Nordisk Foundation under NNF grant number: NNF20CC0035580, NNF Challenge programme CAMiT under Grant agreement: NNF17CO0028232 and the BestTreat project under European Union’s Horizon 2020 research and innovation programme with the Marie Skłodowska-Curie Grant agreement No 813781. We are grateful to Anna-Maria Gutió I Vilardell for assisting in constructing the knockouts, Nicoline Munk Mikkelsen for carrying out some growth characterisations and Carmen Sands for assisting in setting up the flow cytometry experiment and providing (SbU^-^)-GFP. We are also grateful to Tiffany Shang Heng Mak, Carmen Sands and Troels Holger Vaaben for their feedback on the manuscript.

## Author information

### Contributions

K.A.H. and R.V.U. conceived the study. K.A.H. and R.V.U. designed the *in vitro* experiments, and K.A.H, R.V.U. and V.K. designed the *in vivo* experiment. K.A.H. carried out all *in vitro* characterisations, and K.A.H. and V.K. carried out all the *in vivo* characterisations. K.A.H. analysed the data and wrote the manuscript. R.V.U. and M.O.A.S. supervised the study. All authors contributed to the discussion of the results. All authors read and approved the final manuscript.

### Corresponding authors

Correspondence to Ruben Vazquez-Uribe or Morten Otto Alexander Sommer.

### Competing interests

All authors are inventors on a patent filed by DTU on the Biocontainment strategy.

