## Supplementary Information for "Biocontainment strategies for *in vivo* applications of *Saccharomyces boulardii*"

### Supplementary Figures

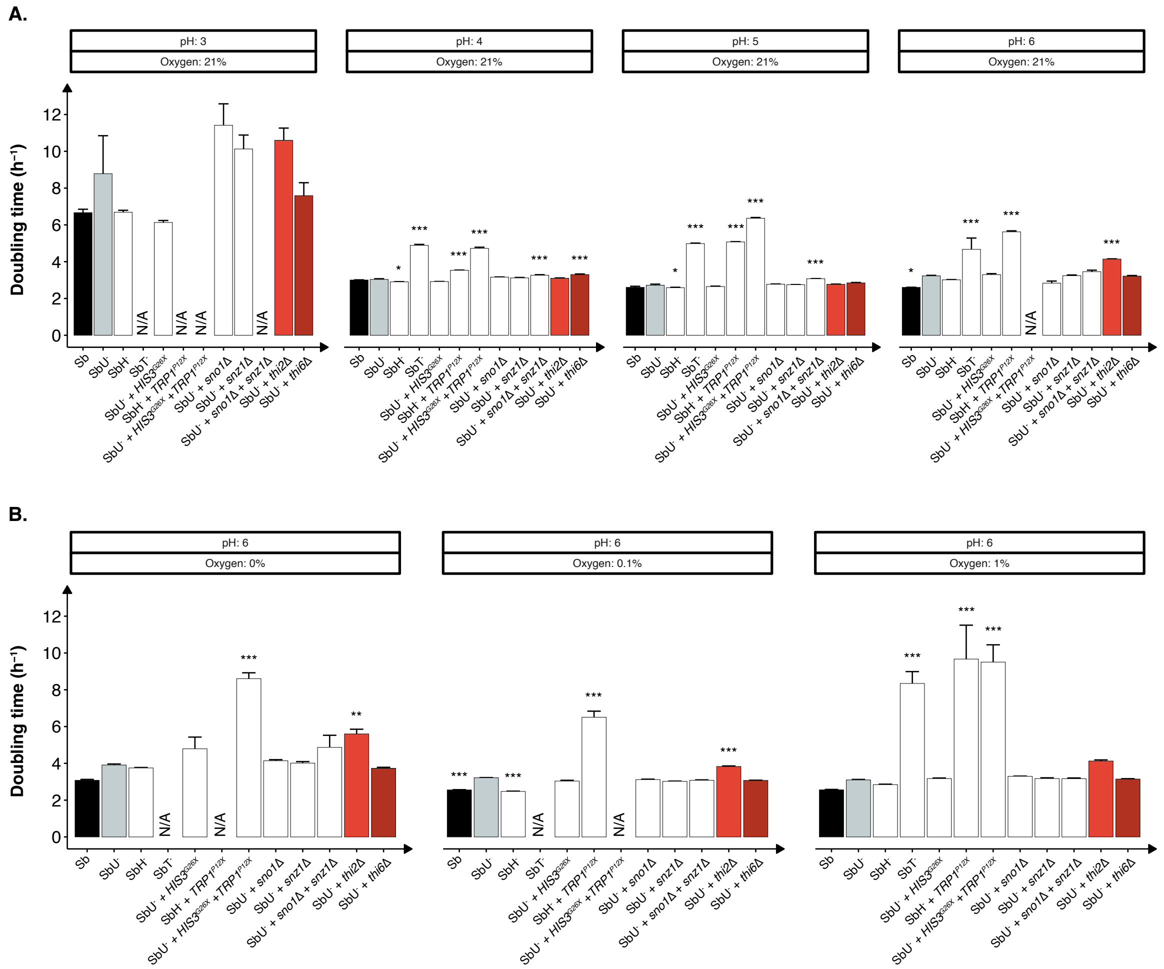


**Figure S1. Growth characterisation of different auxotrophic strains of *S. boulardii*. (A)** Bar plot of the mean doubling time ($h^{-1}$) in pH 3, 4, 5 and 6 at aerobic cultivation. **(B)** Bar plot of the mean doubling time ($h^{-1}$) under anaerobic (0 %) and microaerobic (0.1 % and 1 %) at pH 6. Data presented as mean + SEM (n = 3). One-way ANOVA, Dunnett's post hoc test with Sb *URA3*$^{S81X}$ as reference.

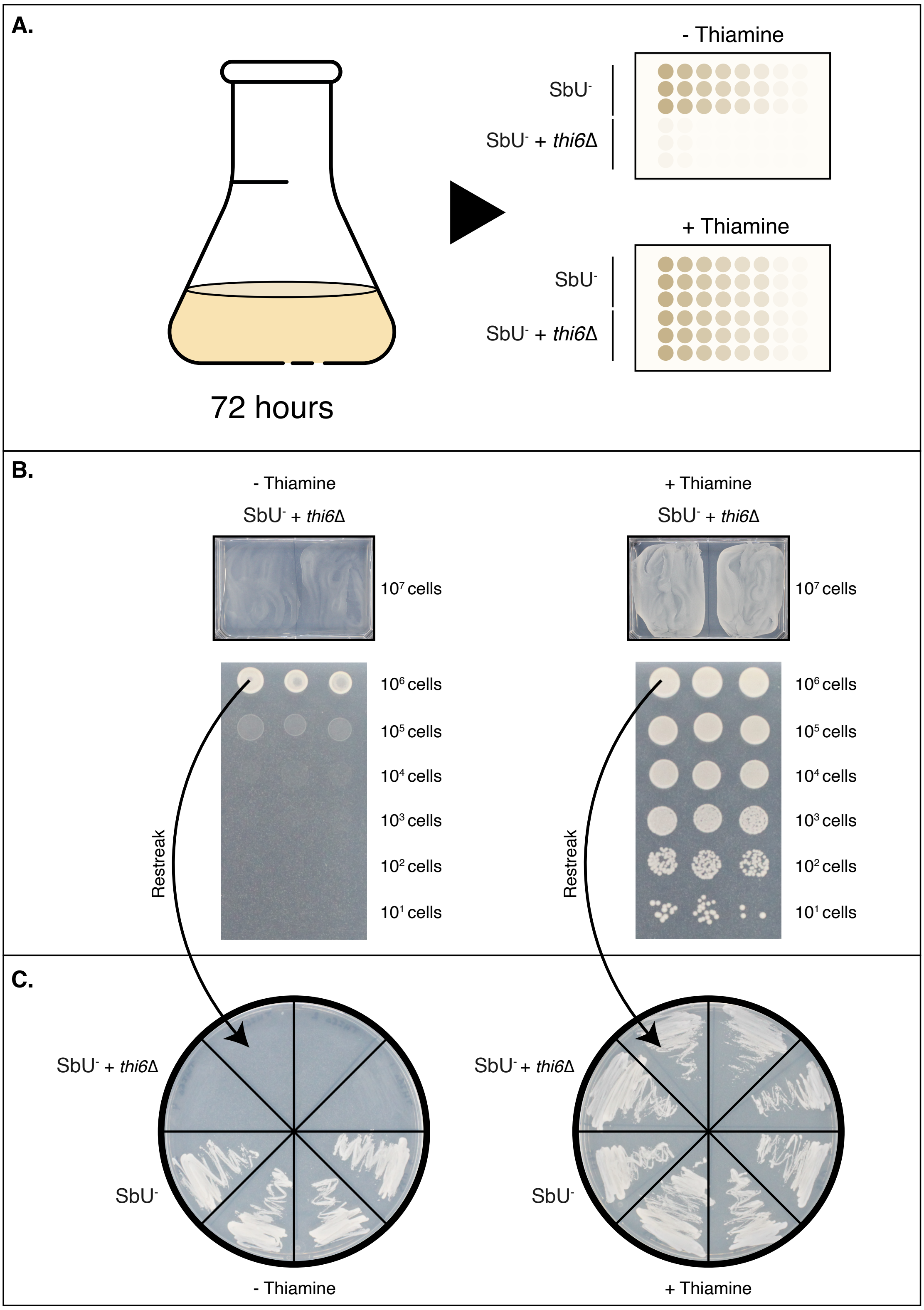


**Figure S2. Escape rate assay of the *thi6*Δ *S. boulardii* strain. (A)** Graphical illustration of the experimental design. The *S. boulardii* strains were cultivated for 72 hours and later spotted on plates with and without thiamine. **(B)** The streaked out and spotted serial dilution of *thi6*Δ *S. boulardii* strain on plates with and without thiamine. **(C)** The grown biomass from the undiluted samples were spread out on fresh selection plates to verify potential escapers.

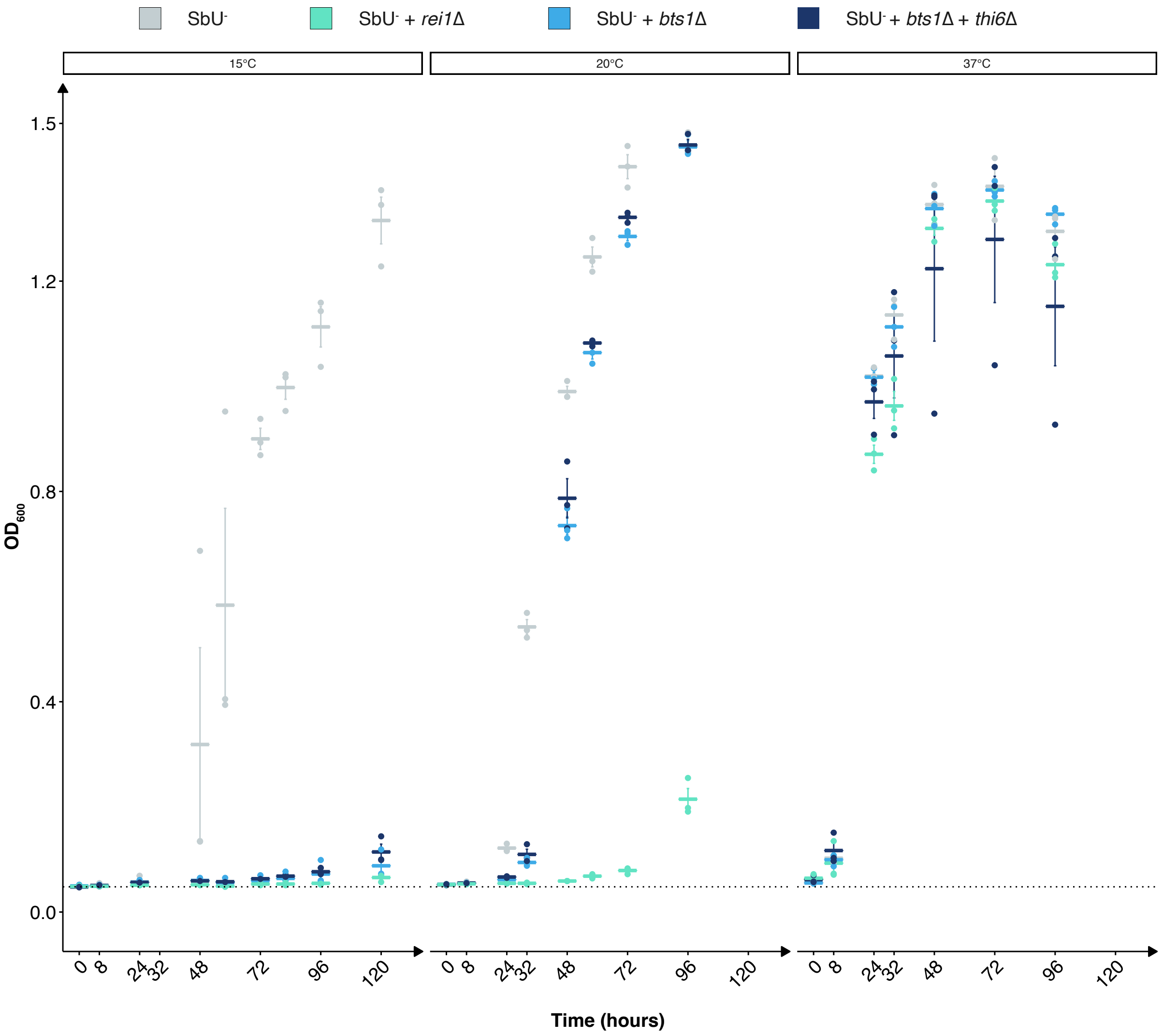


**Figure S3. Growth performance of the cold-sensitive strains at different temperature.** The mean $OD_{600}$ over time at different temperature (15 °C, 20 °C and 37 °C). Data presented as mean ± SEM (n =3). Each point represents a biological replicate.

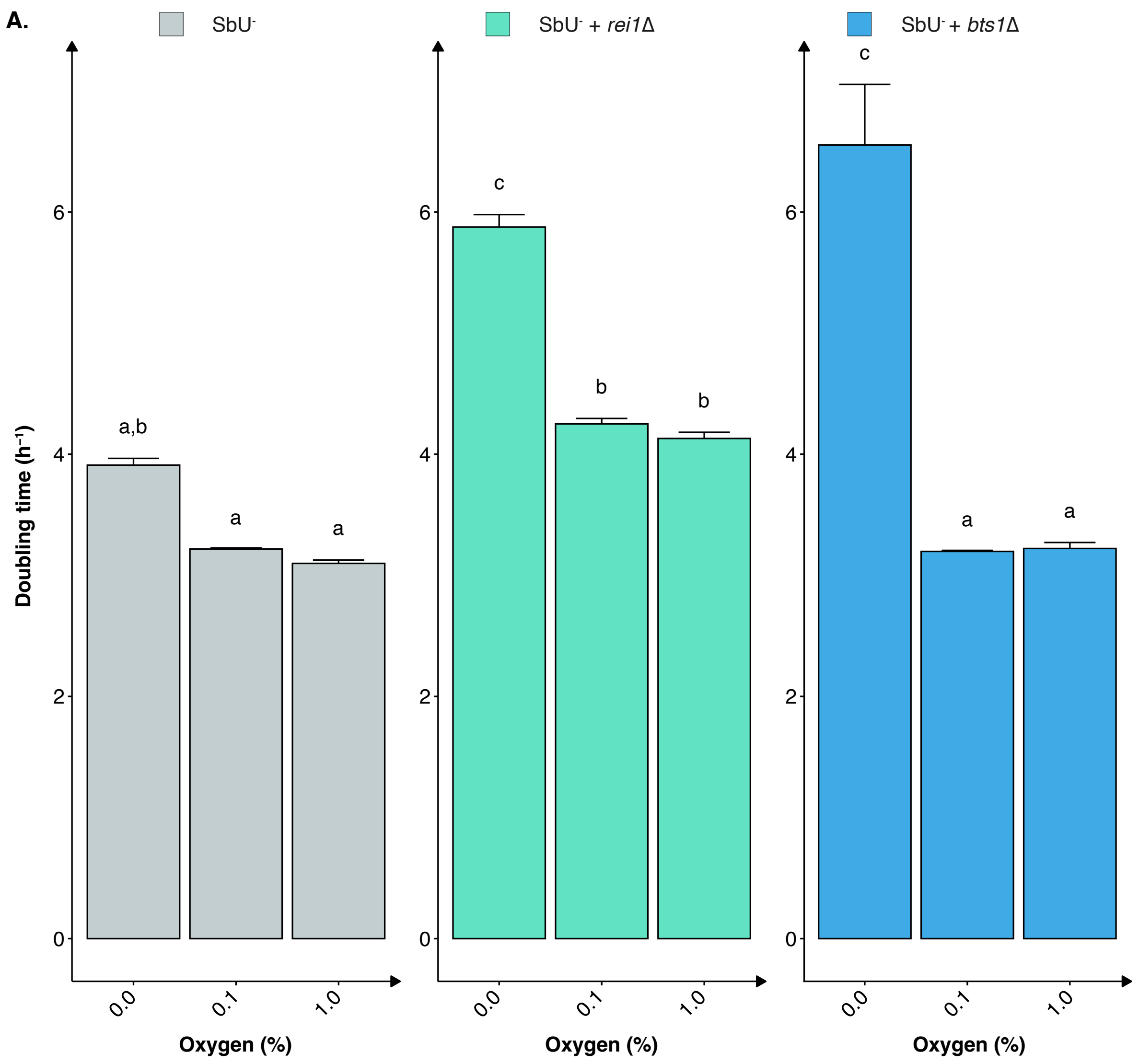


**Figure S4. Growth characterisation of the cold-sensitive strains at different oxygen concentration. (A)** Bar plot of the mean doubling time ($h^{-1}$) under anaerobic (0 %) and microaerobic (0.1 % and 1 %) at pH 6. Data presented as mean + SEM (n = 3). Two-way ANOVA, Tukey post hoc test. The different letters (a, b, and c) above the bars indicate statistically different groups (significance level at $p < 0.05$)

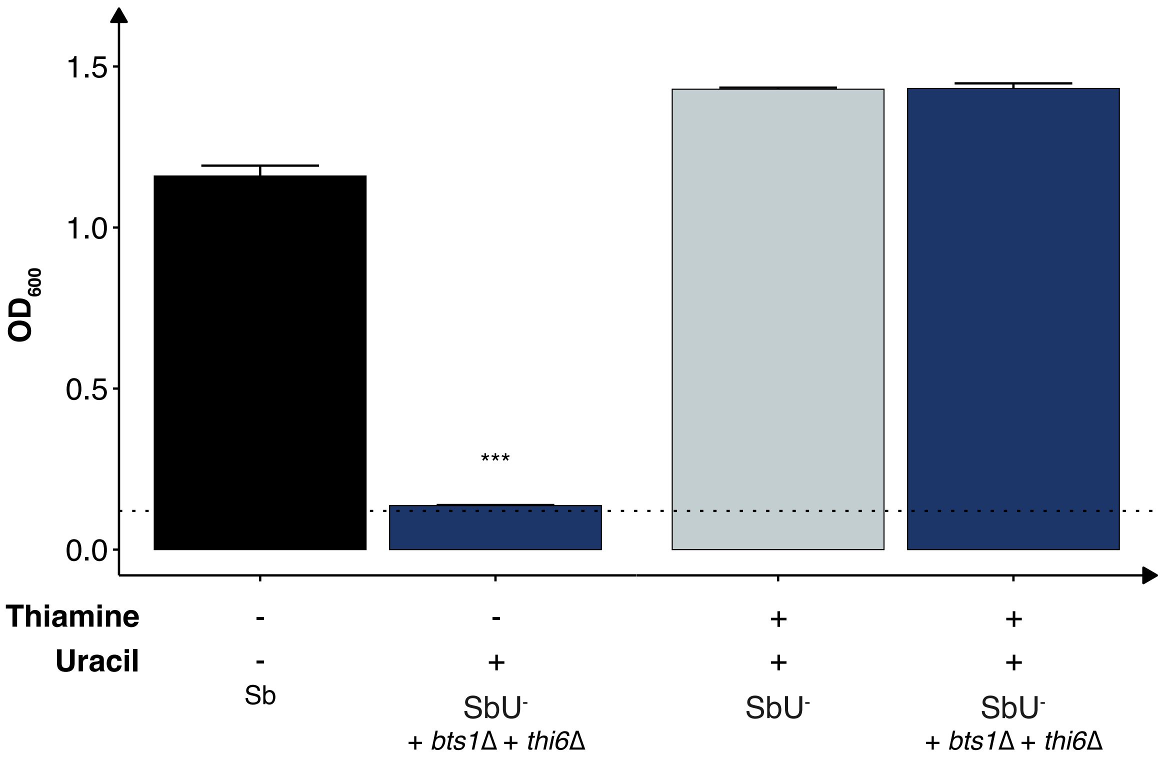


**Figure S5. Characterisation of the combined biocontainment strain.** Bar plot of the mean $OD_{600}$ after 48 hours with (+) and without (-) thiamine supplemented. Data presented as mean + SEM (n = 3). * $p < 0.05$, ** $p < 0.01$ and *** $p < 0.001$. One-way ANOVA, Dunnett's post hoc test with SbU⁻ as reference.

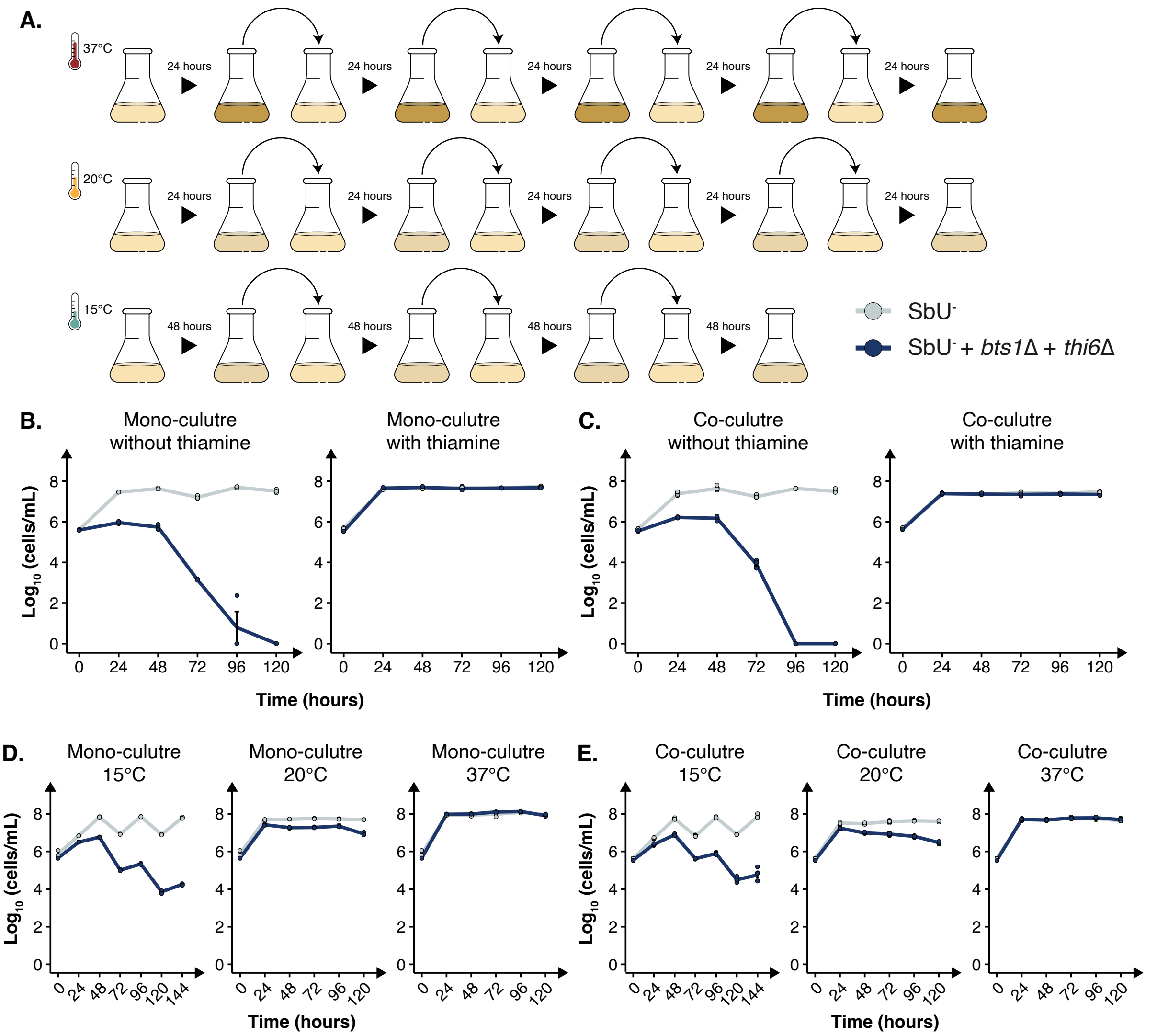


**Figure S6. Growth characterisation of the combined biocontainment strain in mono-culture and co-culture at different condition. (A)** Graphical illustration of the experimental design. SbU$^-$ and SbU$^-$ + *bts1*Δ + *thi6*Δ were inoculated 1:1 in a culture. **(B)** $Log_{10}$ cells/mL of SbU$^-$ and SbU$^-$ + *bts1*Δ + *thi6*Δ in a mono-culture and co-culture with each other in media with and without thiamine. The culture was diluted 1:100 every 48$^{th}$ hour for a total period of 96 hours (n = 3). **(C)** $Log_{10}$ cells/mL of SbU$^-$ and SbU$^-$ + *bts1*Δ + *thi6*Δ in a mono-culture and co-culture with each other at 15°C, 20°C, and 37°C. The culture was diluted 1:100 every 24$^{th}$ hour for a total period of 120 hours for cultures at 20°C and 37°C, and 48$^{th}$ hour for a total period of 144 hours for cultures at 15°C (n = 5). Data presented as mean ± SEM.

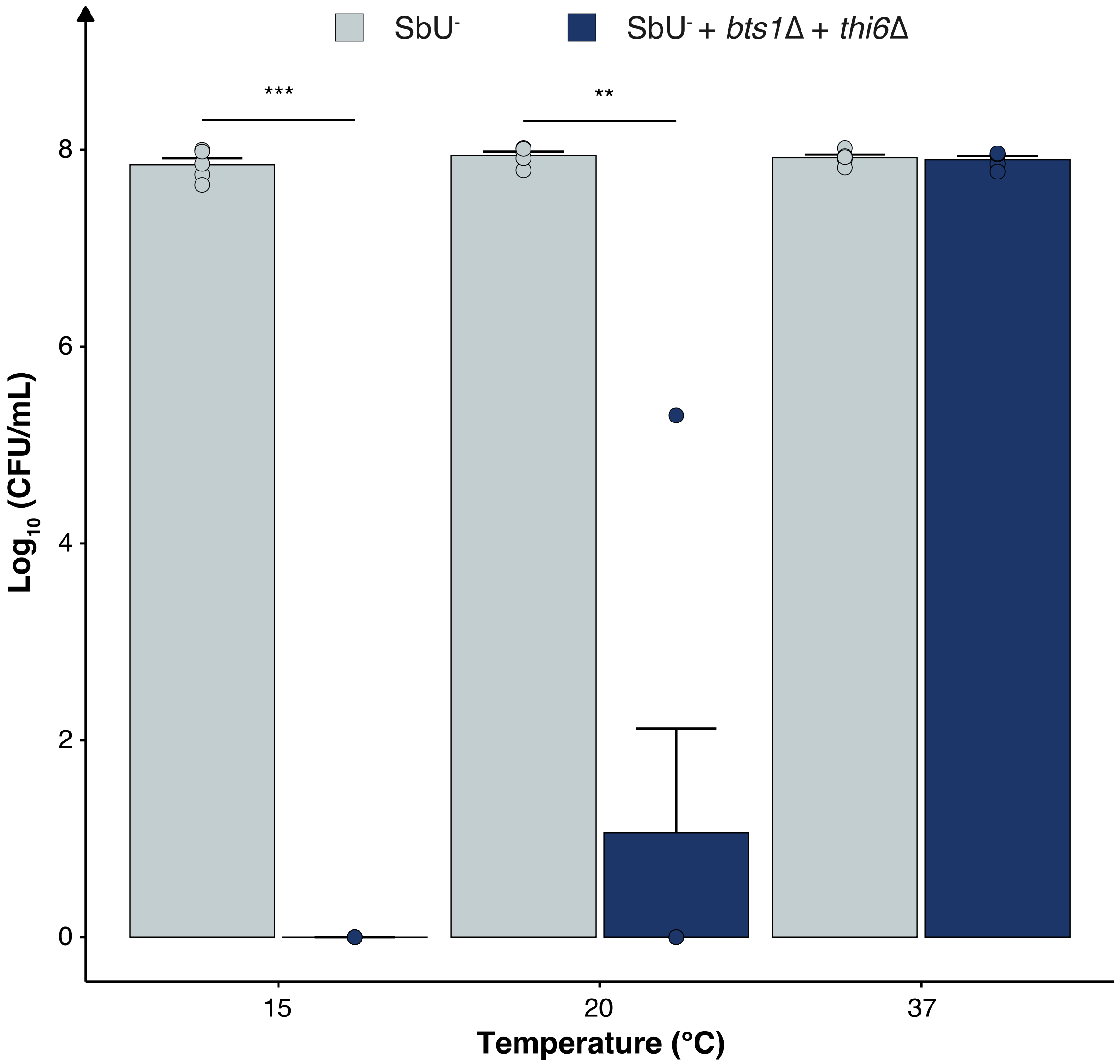


**Figure S7. Survival assay of the combined biocontainment strain.** $Log_{10}$ CFU/mL of $SbU^-$ and $SbU^-$ + *bts1*Δ + *thi6*Δ from a 120 hour co-culture at 37°C that was plated and incubated at 15°C, 20°C, and 37°C for 144, 96 and 48 hours, respectively. Data presented as mean + SEM (n = 5). * $p < 0.05$, ** $p < 0.01$ *** $p < 0.001$. Data was analysed with dependent sample t-test with Bonferroni adjustment for multiple comparisons.

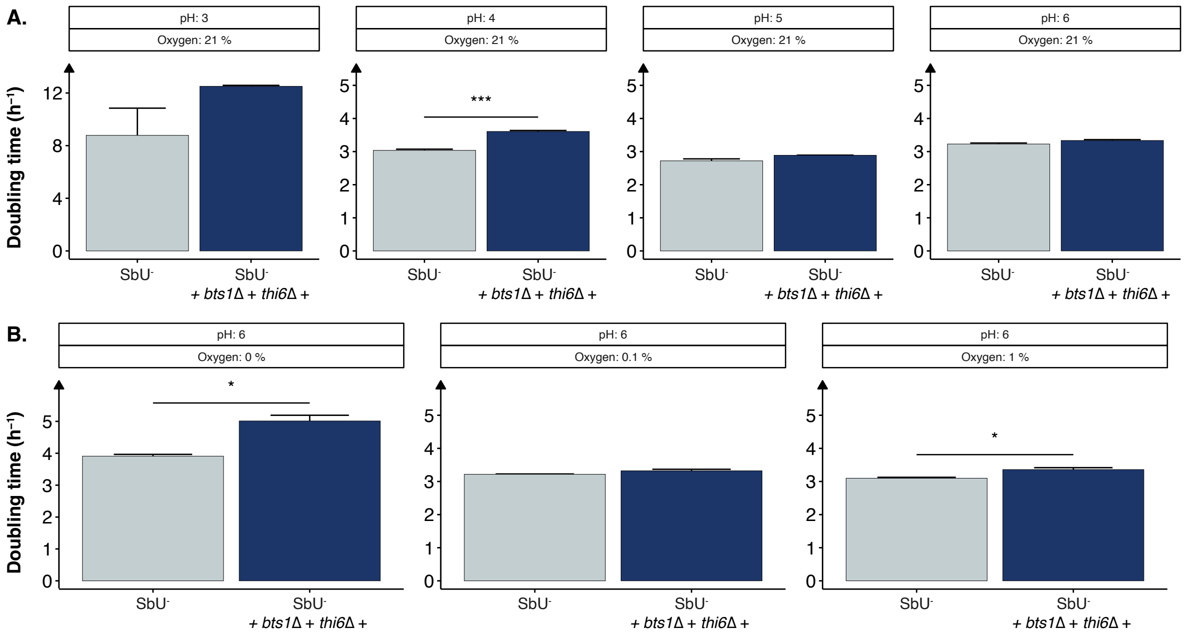


**Figure S8. Growth characterisation of the multi-layered biocontainment strain. (A)** Bar plot of the mean doubling time ($h^{-1}$) in pH 3, 4, 5 and 6 at aerobic cultivation. **(B)** Bar plot of the mean doubling time ($h^{-1}$) under anaerobic (0 %) and microaerobic (0.1 % and 1 %) at pH 6. Data presented as mean + SEM (n = 3). * $p < 0.05$, ** $p < 0.01$ *** $p < 0.001$. All samples analysed with dependent sample t-test.

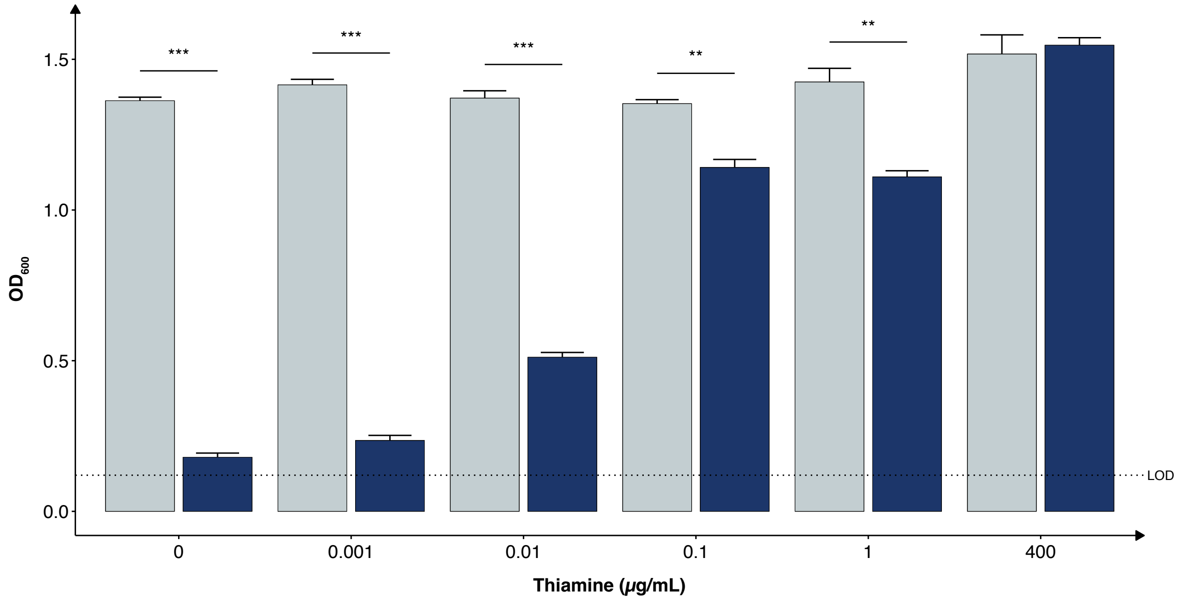


**Figure S9. Growth characterisation of the multi-layered biocontainment strain at different thiamine concentration. (A)** Bar plot of mean $OD_{600}$ after 48 hours under different concentrations of thiamine. Limit of detection (LOD). Data presented as mean + SEM (n = 3). * $p < 0.05$, ** $p < 0.01$ *** $p < 0.001$. All samples analysed with dependent sample t-test.

### Supplementary Tables

#### Table S1. Primers used in this study

| Primer | Sequence | Reference |
|---|---|---|
| pCfB2312-gRNA_fw | atacgaagttatattaagggttg | This study |
| pCfB2312-gRNA_rv | taggcgtatcacgagat | This study |
| pCfB2312_fw | aagcttcagctgacgcgat | This study |
| pCfB2312-rv | tgcaggtcgacaacccttaa | This study |
| URA3repair_fw | tccatggagggcacagttaagccgctaaaggcattataagccaagt acaattttttactc | (Zhang et al., 2014) |
| URA3repair_rv | accaatgtcagcaaattttctgtcttcgaagagtaaaaaattgtacttg gcttataatgc | (Zhang et al., 2014) |
| HIS3repair_fw | gtaaagcgtattacaaatgaaaccaagattcagattgcgatctcttta aagggttaaccc | (Zhang et al., 2014) |
| HIS3repair_rv | ttctgggaagatcgagtgctctatcgctaggggttaaccctttaaaga gatcgcaatctg | (Zhang et al., 2014) |
| TRP1repair_fw | tccgatgctgacttgctgggtattatatgtgtgtaaaatagaaagaga acaattgacccg | (Zhang et al., 2014) |
| TRP1repair_rv | tacaagacttgaaattttccttgcaataaccgggtcaattgttctctttct attttacac | (Zhang et al., 2014) |
| URA3dg-fw | cttgcatgacaattctgcta | This study |
| URA3dg-rv | ttcttaacccaactgcacag | This study |
| HIS3dg-fw | aaagatctaccaccgctctg | This study |
| HIS3dg-rv | gcgattggcattatcacata | This study |
| TRP1dg-fw | gccgattaagaattcggtcg | This study |
| TRP1dg-rv | gcactgagtagtatgttgcag | This study |
| BTS1dg_fw | aaatcgcgaatttaccggcg | This study |
| BTS1dg_rv | gccgccatctctactcactc | This study |
| BTS1repair_fw | tcattttcaaagaagctactaatagaaagagaacaaagcgtttacga gtctggaaaatcaaataaattgatcaatcaaattagtggaggaagat agtcagaaataaagccttctctcctc | This study |
| BTS1repair_rv | gaggagagaaggctttatttctgactatcttcctccactaatttgattga tcaatttatttgattttccagactcgtaaacgctttgttctctttctattagt agcttctttgaaaatga | This study |
| REI1dg_fw | ttgattcgctgcctgttcg | This study |
| REI1dg_rv | cccgatattccccgtgactc | This study |
| REI1repair_fw | tacaggttatgagatgcttctcattagaagtcaagaagagagcatatc agtaacaatacgttctttttgcactactttttttagtattttgtcgcatatat actgcttcaccatttgtac | This study |

| | | |
|---|---|---|
| REI1repair_rv | gtacaaatggtgaagcagtatatatgcgacaaaatactaaaaaaagtagtgcaaaaagaacgtattgttactgatatgctctcttcttgacttctaatgagaagcatctcataacctgta | This study |
| SNZ1dg_fw | agctttaccctggaagcacc | This study |
| SNZ1dg_rv | ggacgctgagagctatggac | This study |
| SNZ1repair_fw | agaaacctttaggaacgactagcaaatatacacagtactaatattcagttaattatcacgtttctttgaacaggtattttgagcattataacactttttcccccctcaactttgtattac | This study |
| SNZ1repair_rv | gtaatacaaagttgagggggaaaaagtgttataatgctcaaaatacctgttcaaagaaacgtgataattaactgaatattagtactgtgtatatttgctagtcgttcctaaaggtttct | This study |
| SNO1dg_fw | ccttacccttgtcggctgag | This study |
| SNO1dg_rv | aagggcctgcggaagatcac | This study |
| SNO1repair_fw | aggtttttttttcttattatttcatttcgttaaatagaaagaaaaaccatatcttaaagtaataccgccggttttccacatttttatattaccaaaacctgagagatttttttcacatcga | This study |
| SNO1repair_rv | tcgatgtgaaaaaatctctcaggttttggtaatataaaaatgtggaaaaccggcggtattactttaagatatggtttttctttctatttaacgaaatgaaataataagaaaaaaaaacct | This study |
| THI2repair_fw | ttttcatttcatttcaccacgtatatatatagcctatatatatatccgcactagaaccaagcttattgagccttcccttcactcatttcaagaaaaaaaaagccaaaagctttgcttgga | This study |
| THI2repair_ rv | tccaagcaaagcttttggcttttttttcttgaaatgagtgaagggaaggctcaataagcttggttctagtgcggatatatatataggctatatatatacgtggtgaaatgaaatgaaaa | This study |
| THI2_dg fw | tacaatggcagccctcttgg | This study |
| THI2_dg rv | ccctggcagataggaaaccc | This study |
| THI6repair_fw | aaccatggttctcaagaccaaatcactctgaagctaaattatttaaatcaaacagcggaaactatttacaacgtaatgattttataaactgattaaataaggaagaaagcccaaaaaatt | This study |
| THI6repair_rv | aattttttgggctttcttccttatttaatcagtttataaaatcattacgttgtaaatagtttccgctgtttgatttaaataatttagcttcagagtgatttggtcttgagaaccatggtt | This study |
| THI6_dg fw | tccgtttctagctgcaggtc | This study |
| THI6_dg rv | agtttgttcctcctggggtg | This study |
| pCfB3050_fw | cttcgctattacgccagctg | This study |
| pCfB3050_rv | ccgattcattaatgcagctg | This study |

**Table S2. Plasmids used in this study**

| Plasmid name | Genotype | Marker (*E. coli / S. boulardii*) | Reference |
|---|---|---|---|
| pCfB2312 | CEN6_ARS4 Amp $P_{TEF1}$-Cas9-$t_{CYC1}$ kanMX | Amp / KanMX | (Jessop-Fabre et al., 2016) |
| pCfB2312-URA3 | pCfB2312; $P_{SNR52}$-gRNA(*URA3*)-$t_{SUP4}$ | Amp / KanMX | This study |
| pCfB2312-HIS3 | pCfB2312; $P_{SNR52}$-gRNA(*HIS3*)-$t_{SUP4}$ | Amp / KanMX | This study |
| pCfB2312-TRP1 | pCfB2312; $P_{SNR52}$-gRNA(*TRP1*)-$t_{SUP4}$ | Amp / KanMX | This study |
| pCfB2909-Exe4 | pCfB2909: $P_{TDH3}$-Exe4-$t_{DIT1**}$ | Amp | (Hedin et al., 2022) |
| pCfB2909-GFP | pCfB2909: $P_{TDH3}$-GFP- $t_{ADH1}$ | Amp | This study |
| pCfB2909-mKate | pCfB2909: $P_{TDH3}$-mKate-$t_{ADH1}$ | Amp | This study |
| pCfB3050 (URA3) | pESC; $P_{SNR52}$-gRNA(XII-5)-$t_{SUP4}$ | Amp / *URA3* | This study |
| pCfB3050-THI2 | pESC; $P_{SNR52}$-gRNA(*THI2*)-$t_{SUP4}$ | Amp / *URA3* | This study |
| pCfB3050-THI6 | pESC; $P_{SNR52}$-gRNA(*THI6*)-$t_{SUP4}$ | Amp / *URA3* | This study |
| pCfB3050-SNO1 | pESC; $P_{SNR52}$-gRNA(*SNO1*)-$t_{SUP4}$ | Amp / *URA3* | This study |
| pCfB3050-SNZ1 | pESC; $P_{SNR52}$-gRNA(*SNZ1*)-$t_{SUP4}$ | Amp / *URA3* | This study |
| pCfB3050-BTS1 | pESC; $P_{SNR52}$-gRNA(*BTS1*)-$t_{SUP4}$ | Amp / *URA3* | This study |
| pCfB3050-REI1 | pESC; $P_{SNR52}$-gRNA(*REI1*)-$t_{SUP4}$ | Amp / *URA3* | This study |

**Table S3. Construct sequences used in this study**

| Name | Sequence | Reference(s) |
|---|---|---|
| P$_{TDH3}$ | cagttcgagtttatcattatcaatactgccatttcaaagaatacgtaaataattaa<br>tagtagtgattttcctaactttatttagtcaaaaaattggccttttaattctgctgta<br>acccgtacatgcccaaaatagggggcgggttacacagaatatataacatcat<br>aggtgtctgggtgaacagtttattcctggcatccactaaatataatggagccc<br>gctttttaagctggcatccagaaaaaaaaagaatcccagcaccaaaatattgt<br>tttcttcaccaaccatcagttcataggtccattctcttagcgcaactacacagaa<br>caggggcacaaacaggcaaaaaacgggcacaacctcaatggagtgatgc<br>aacctgcttggagtaaatgatgacacaaggcaattgacctacgcatgtatcta<br>tctcattttcttacaccttctattaccttctgctctctctgatttggaaaaagctgaa<br>aaaaaaggttgaaaccagttccctgaaattattcccctatttgactaataagtat<br>ataaagacggtaggtattgattgtaattctgtaaatctatttcttaaacttcttaaa<br>ttctacttttatagttagtcttttttttagttttaaaacactaagaacttagtttcgaat<br>aaacacacataaacaaacaaa | This study |
| t$_{ADH1}$ | gtagatacgttgttgacacttctaaataagcgaatttcttatgatttatgatttttat<br>tattaaataagttataaaaaaaataagtgtatacaaattttaaagtgactcttagg<br>ttttaaaacgaaaattcttattcttgagtaactctttcctgtaggtcaggttgctttc<br>tcaggtatagcatgaggtcgctc | This study |
| mKate | atggtttctgaactcatcaaggaaaacatgcacatgaaactttacatggaagg<br>tactgtgaacaatcatcattttaagtgtacatccgagggtgaaggcaaacctta<br>cgaaggaactcaaactatgagaattaaagctgtagaaggtggaccattacct<br>tttgcatttgatatcttggcaacatcattcatgtatgggagcaagacattcataa<br>accatactcaaggtataccagactttttcaaacagagttttccagagggttttac<br>atgggaaagagtaacaacgtacgaggatggaggtgtattgacagccactca<br>agacacatcacttcaagatgggtgtttaatctacaatgtcaagattagaggcg<br>tcaatttcccttctaatggtccagttatgcagaaaaagacattaggctgggaa<br>gcgtcaaccgaaaccctgtaccctgctgatggtggcctagaaggcagagct<br>gacatggcccttaaactggttggtggagggcatctaatctgcaatttgaaaac<br>cacttatcgttctaaaaagccagccaaaaacctaaagatgccaggtgtttact<br>acgtcgaccgaagattagaaaggattaaagaggctgataaagagacttatgt<br>tgaacaacacgaagtggcagtggctagatactgtgatttgccatctaagttgg<br>gacacagataa | This study |
| yEGFP | atgtctaaaggtgaagaattattcactggtgttgtcccaattttggttgaattaga<br>tggtgatgttaatggtcacaaattttctgtctccggtgaaggtgaaggtgatgc<br>tacttacggtaaattgaccttaaaatttatttgtactactggtaaattgccagttcc<br>atggccaaccttagtcactactttcggttatggtgttcaatgttttgcgagatac<br>ccagatcatatgaaacaacatgactttttcaagtctgccatgccagaaggttat<br>gttcaagaaagaactatttttttcaaagatgacggtaactacaagaccagagc<br>tgaagtcaagtttgaaggtgataccttagttaatagaatcgaattaaaaggtatt<br>gattttaaagaagatggtaacattttaggtcacaaattggaatacaactataact<br>ctcacaatgtttacatcatggctgacaaacaaaagaatggtatcaaagttaact<br>tcaaaattagacacaacattgaagatggttctgttcaattagctgaccattatca<br>acaaaatactccaattggtgatggtccagtcttgttaccagacaaccattactt<br>atccactcaatctgccttatccaaagatccaaacgaaaagagagaccacatg<br>gtcttgttagaatttgttactgctgctggtattacccatggtatggatgaattgta<br>caaatga | This study |

| | | |
|---|---|---|
| Exendin-4 | tctaccaacggaatgcgtgcgatcgcgtgcattccgagtttatcattatcaata<br>ctgccatttcaaagaatacgtaaataattaatagtagtgattttcctaactttattt<br>agtcaaaaaattagccttttaattctgctgtaacccgtacatgcccaaaatagg<br>gggcgggttacacagaatatataacatcgtaggtgtctgggtgaacagtttat<br>tcctggcatccactaaatataatggagcccgctttttaagctggcatccagaa<br>aaaaaaagaatcccagcaccaaaatattgttttcttcaccaaccatcagttcat<br>aggtccattctcttagcgcaactacagagaacaggggcacaaacaggcaa<br>aaaacgggcacaacctcaatggagtgatgcaacctgcctggagtaaatgat<br>gacacaaggcaattgacccacgcatgtatctatctcattttcttacaccttctatt<br>accttctgctctctctgatttggaaaaagctgaaaaaaaaggttgaaaccagtt<br>ccctgaaattattcccctacttgactaataagtatataaagacggtaggtattga<br>ttgtaattctgtaaatctatttcttaaacttcttaaattctacttttatagttagtcttttt<br>tttagttttaaaacaccaagaacttagtttcgaataaacacacataaacaaaca<br>aaaacaaaatgagatttccatctatttttactgctgttttgtttgctgcttcttctgc<br>tttggctgctccagttaatactactactgaagatgaaactgctcaaattccagct<br>gaagctgttattggttattctgatttggagggtgactttgatgttgctgttttgcca<br>ttttctaactctactaacaacggtttgctattcatcaacactactatcgcttctatc<br>gctgctaaagaagaaggtgtttctttggataaaagagaagaaggtgaaccaa<br>aacatggtgaaggcacattcacatctgatctgtccaaacaaatggaggagg<br>aagcggtacgtttatttattgaatggttaaaaaacgggggacctagctccggc<br>gcgcccccccccgagctaataaagtaagagcgctacattggtctacctttttctt<br>ttacttaaacattagttagttcgttttctttttctttttttatgtttccccccccaaagttc<br>tgattttataatattttatttcacacaattccatttaacagagggggaatagattct<br>ttagcttagaaaattagtgatcaatatatatttgcctttcttttcatcttttcagtgat<br>attaatggtttcgagacactgcaatggccctactagtgctgaggcattaat | (Hedin et al., 2022) |

**Table S4. gRNA used in this study**

| gRNA | Sequence | Reference |
|---|---|---|
| URA3 | gagtaaaaaattgtacttgg | (Zhang et al., 2014) |
| HIS3 | ccctttaaagagatcgcaat | (Zhang et al., 2014) |
| TRP1 | gtcaattgttctctttctat | (Zhang et al., 2014) |
| THI2 | actacaattatctccatgtt | This study |
| THI6 | ttaaataaccataaaatgaa | This study |
| SNO1 | gacgccttaattattcccgg | This study |
| SNZ1 | acccaactgcattaacaatg | This study |
| BTS1 | ttgctgaggacattacagag | This study |
| REI1 | gaagatgactgggaagacgt | This study |

**Table S5. Strains used in this study**

| Strain | Genotype | Marker | Parental strain | Reference |
|---|---|---|---|---|
| Sb | | N/A | SB-ATCC-796 | This study |
| $SbU^-$ | Sb $URA3^{S81X}$ | N/A | Sb | This study |
| $SbH^-$ | Sb $HIS3^{G26X}$ | N/A | Sb | This study |
| $SbT^-$ | Sb $TRP1^{P12X}$ | N/A | Sb | This study |
| $SbU^-$ + $HIS3^{G26X}$ | Sb $URA3^{S81X}$ + $HIS3^{G26X}$ | N/A | $SbU^-$ | This study |
| $SbH^-$ + $TRP1^{P12X}$ | Sb $HIS3^{G26X}$ + $TRP1^{P12X}$ | N/A | $SbH^-$ | This study |
| $SbU^-$ + $HIS3^{G26X}$ + $TRP1^{P12X}$ | Sb $URA3^{S81X}$ + $HIS3^{G26X}$ | N/A | $SbU^-$ + $HIS3^{G26X}$ | This study |
| $SbU^-$ + *thi6*Δ | Sb $URA3^{S81X}$ + *thi6*Δ | N/A | $SbU^-$ | This study |
| $SbU^-$ + *thi2*Δ | Sb $URA3^{S81X}$ + *thi2*Δ | N/A | $SbU^-$ | This study |
| $SbU^-$ + *sno1*Δ | Sb $URA3^{S81X}$ + *sno1*Δ | N/A | $SbU^-$ | This study |
| $SbU^-$ + *snz1*Δ | Sb $URA3^{S81X}$ + *snz1*Δ | N/A | $SbU^-$ | This study |
| $SbU^-$ + *sno1*Δ + *snz1*Δ | Sb $URA3^{S81X}$ + *sno1*Δ + *snz1*Δ | N/A | $SbU^-$+ *sno1*Δ | This study |
| $SbU^-$ + *rei1*Δ | Sb $URA3^{S81X}$ + *rei1*Δ | N/A | $SbU^-$ | This study |
| $SbU^-$ + *bts1*Δ | Sb $URA3^{S81X}$ + *bts1*Δ | N/A | $SbU^-$ | This study |
| $SbU^-$ + *bts1*Δ + *thi6*Δ | Sb $URA3^{S81X}$ + *bts1*Δ + *thi6*Δ | N/A | $SbU^-$+ *bts1*Δ | This study |
| ($SbU^-$)-Exe4 | $SbU^-$ + XII-5 $P_{TDH3}$-Exe4-$t_{DIT1}$** | N/A | $SbU^-$ | (Hedin et al., 2022) |
| ($SbU^-$ + *thi6*Δ)-Exe4 | $SbU^-$ + *thi6*Δ + XII-5 $P_{TDH3}$-Exe4-$t_{DIT1}$** | N/A | $SbU^-$ + *thi6*Δ | This study |
| ($SbU^-$ + *bts1*Δ)-Exe4 | $SbU^-$ + *bts1*Δ + XII-5 $P_{TDH3}$-Exe4-$t_{DIT1}$** | N/A | $SbU^-$+ *bts1*Δ | This study |
| ($SbU^-$ + *bts1*Δ + *thi6*Δ)-Exe4 | $SbU^-$ + *bts1*Δ + *thi6*Δ + XII-5 $P_{TDH3}$-Exe4-$t_{DIT1}$** | N/A | $SbU^-$ + *bts1*Δ + *thi6*Δ | This study |

| | | | | |
|---|---|---|---|---|
| (SbU$^-$)-GFP | SbU$^-$ + XII-5 $P_{TDH3}$-yEGFP-$t_{ADH1}$ | N/A | SbU$^-$ | This study |
| (SbU$^-$ + *bts1*Δ + *thi6*Δ)-mKate | SbU$^-$ + *bts1*Δ + *thi6*Δ + XII-5 $P_{TDH3}$-mKate-$t_{ADH1}$ | N/A | SbU$^-$ + *bts1*Δ + *thi6*Δ | This study |
